# DPCfam: a new method for unsupervised protein family classification

**DOI:** 10.1101/2020.07.30.224592

**Authors:** Elena Tea Russo, Alessandro Laio, Marco Punta

## Abstract

**Motivation:** As the UniProt database approaches the 200 million entries’ mark, the vast majority of proteins it contains lack any experimental validation of their functions. In this context, the identification of homologous relationships between proteins remains the single most widely applicable tool for generating functional and structural hypotheses in silico. Although many databases exist that classify proteins and protein domains into homologous families, large sections of the sequence space remain unassigned.

**Results:** We introduce DPCfam, a new unsupervised procedure that uses sequence alignments and Density Peak Clustering to automatically classify homologous protein regions. Here, we present a proof-of-principle experiment based on the analysis of two clans from the Pfam protein family database. Our tests indicate that DPCfam automatically-generated clusters are generally evolutionary accurate corresponding to one or more Pfam families and that they cover a significant fraction of known homologs. Overall, DPCfam shows potential both for assisting manual annotation efforts (domain discovery, detection of classification inconsistencies, improvement of family coverage and boosting of clan membership) and as a stand-alone tool for unsupervised classification of sparsely annotated protein datasets such as those from environmental metagenomics studies (domain discovery, analysis of domain diversity).

**Availability:** Algorithm implementation used in this paper is available at https://gitlab.com/ETRu/dpcfam (Requires Python 3, C++ compiler and runs on Linux systems.); data are available at https://zenodo.org/record/3934399

## 1 Introduction

Conserved evolutionary modules shared by different proteins typically present some degree of structural and, to a lesser extent, functional similarity [30]. The identification of homologous protein families is thus of great importance for in silico protein annotation. Several resources have been developed toward this goal, including but not limited to, Pfam [10], SMART [20], TIGRFAMs [13], PANTHER [25], SFLD [1], CATH-Gene3D [21], SUPERFAMILY [36] and ECOD [7]. Some databases restrict themselves to specific functional categories (SMART, SFLD), phylogenetic groups (TIGRFAMs) or to families for which structural information is available (CATH-Gene3D, SUPERFAMILY, ECOD). Others aim to classify the protein sequence space more widely (Pfam, PANTHER). Most databases try to identify domains (evolutionary, structural and/or functional units) while some build families for full-length protein sequences (TIGRFAM, PANTHER). All of these resources take advantage, at some level, of expert manual curation. While this helps increasing quality of defined families, it can limit the area of the sequence space that each classification is able to cover at any point in time. For example, Pfam residue coverage of the UniProtKB database as of release 04/2018 [10] was around 53% with more than 20% of all UniProtKB sequences lacking any type of Pfam annotation. In order to alleviate this problem, databases have been developed that integrate into a single platform families from several other resources (InterPro [28], CDD [24]).

An alternative approach to manually curated family classification is performing automatic, sequence-based clustering of protein regions. Although unsupervised clustering will generally result in a less accurate classification, it has the advantage that high coverage of the protein sequence space can be achieved, at limited cost. As a consequence, it can be used effectively to complement manual annotations. Automated family classification has a long history in protein bioinformatics and over the years has led to the development of algorithms such as ADDA [14], COG [35], EVEREST [31], and MCL [11], among others. Until 2015, the ADDA clustering algorithm was used to produce Pfam-B, which was an automatically-built companion to the manually curated Pfam main family collection. Pfam-B, however, has since been discontinued due to the heavy computational cost of running ADDA on modern day, large sequence databases [12]. MCL [11] has been widely used for automatic clustering of both amino-acid and nucleotide sequences (e.g. [8], [27], [3], [23], [19]). MCL underlying algorithm groups protein sequences into families using stochastic flows on pre-computed graphs where nodes represent protein sequences and edges are sequence similarity scores obtained from pairwise alignments. A recent high-performance parallel implementation of MCL (HIpMCL) allows for efficient clustering of large-scale networks [2]. Although MCL is very effective, it is not meant to cluster together individual domains but rather full-length sequences in this being similar to COG [35] and other ortholog/paralog clustering methods. Here we introduce DPCfam, a new method for automated classification of homologous protein sequences into families based on all-versus-all sequence similarity scores and Density Peak Clustering (DPC) [33]. Given an initial list of query sequences and a dataset of search sequences (the latter including the queries), we run BLAST alignments of all query vs all search sequences. Next, search sequence regions that align to similar parts of a query are grouped together into primary clusters by DPC. Primary clusters are grouped into “metaclusters” (MCs) based on the number of search sequence regions they have in common and on the correspondence between their respective boundaries. Finally, MCs are merged if they still share a significant number of search sequence regions. The sequences that form an MC can be used as seeds to build a Multiple Sequence Alignment (MSA) and a corresponding profile-HMM, similarly to what is done in Pfam [10]. In this work, we use the Pfam database family classification as our ‘ground truth’ for protein annotation. In particular, by analyzing two Pfam clans (also known as superfamilies) we provide a proof-of-principle of the ability of DPCfam to produce MCs that are constituted of protein sequences sharing a core homologous region (families). We show that the families produced by DPCfam can represent single domains or, sometimes, conserved domain architectures. Although in principle the DPCfam algorithm can be applied to the analysis of large sequence databases such as UniprotKB or metagenomic datasets, the specific implementation presented in this paper is only suitable for analysis of relatively small datasets. We are currently working on a parallelized c++ version of the algorithm which, when ran on Tier-0 supercomputers, we estimate will take about 50,000 CPU hours to complete the clustering of the whole UniRef50 database (v 07/2017, about 23 million sequences).

## 2 Methods

In the following sections we describe the DPCfam workflow, which consists of: running BLAST alignments of our query (clan) database against Uniref50 (2.1), primary clustering of alignments falling on the same query sequence (2.2.1), metaclustering of primary clusters (2.2.2) and, finally, merging of metaclusters (2.2.3). In sections 2.3 and 3.4 we describe our protocols for comparing DPCfam clusters with the Pfam-annotated families. Finally, in section 2.5, we report on the choice of the two Pfam clans that we analyse here.

### 2.1 BLAST searches

Our database of reference throughout this work is Uniref50 (v. 07/2017). Given a Pfam clan, for example PUA, we generate a dataset constituted of all Uniref50 full-length sequences that carry a PUA clan member annotation by matching their UniProtKB ids with those of sequences in Pfam-A.full v.31. We name such a dataset PUA_UR50. Next, for each sequence (query) in the dataset, we perform a local alignment search against the full Uniref50 database using NCBI BLAST (v. 2.2.30+)[6] and save all alignments with E-value < 0.1 (up to 5 millions, using the max_target_seqs option of BLAST).

We define a *BLAST alignment*, labeled by an index *i*, as:

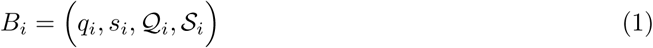

where *q*_*i*_ is the identifier of the query sequence, *s*_*i*_ is the identifier of the search sequence, *𝒬*_*i*_ and *𝒮*_*i*_ are regions on, respectively, the query and the search sequence. More in detail, *𝒬*_*i*_ and *𝒮*_*i*_ represent the boundaries (start and end points) of the pairwise alignments on the query search sequences, respectively (see Figure S1 A). Note that gaps and insertions are not taken into account.

### 2.2 Clustering of BLAST alignments

DPC [33] entails the following steps: i) defining a distance in the space of the objects that are to be clustered; ii) for each object estimating its local density using the distance defined in (i); iii) selecting density peaks (cluster centers) and, finally, iv) assigning of non-peak objects to density peaks (clustering). In DPCfam we perform two rounds of DPC. The first round allows us clustering alignments that cover similar regions of the query sequences (primary clusters); in the second round we group together primary clusters that share a number of overlapping alignments (metaclusters). As a final step, in DPCfam, we apply an *ad hoc* merging procedure to further group metaclusters that share many of their member alignments. Alignments belonging to merged metaclusters can then be linked back to the respective aligned sequences, thus obtaining clusters of protein regions, which are meant to represent families (see Results).

#### 2.2.1 Primary clustering

For a query *q*_0_ we write the set of all of its alignments as:

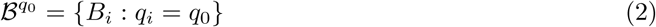

We define the distance between alignments in 𝔅^*q*0^ as:

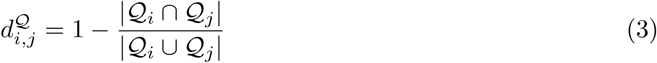

where |*𝒬*_*i*_ ⋂ *𝒬*_*j*_| is the length (intended as number of residues) of the intersection between the segments identified by *𝒬*_*i*_ and *𝒬*_*j*_, while |*𝒬*_*i*_ ⋃ *𝒬*_*j*_| is the length of their union (see Figure S1 B). This distance is 0 if *B*_*i*_ and *B*_*j*_ are aligned to the same portion of the query *q*_0_, that is, *𝒬* _*i*_ = *𝒬*_*j*_; while it is 1 if *𝒬*_*i*_ and *𝒬*_*j*_ do not overlap at all. As defined, 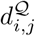 represents a metric since it is symmetric and satisfies the triangular inequality. Using the distance in Eq. 3, we estimate the density *ρ*_*i*_ of the alignment *i*:

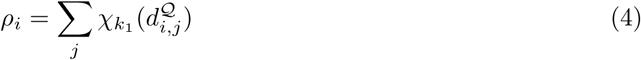

where *𝒳k*_1_ (*x*) = 1 if *x< k*_1_ and zero otherwise. Thus, the density of an alignment *B*_*i*_ is given by the number of alignments that belong to the same set 𝔅^*q*0^ and that are found at a distance less than *k*_1_ from *B*_*i*_. In the algorithm, we set *k*_1_ = 0.2, according to the rule of thumb in [33]: the average number of neighbours closer than *k*_1_ to a point is around 1 to 2% of the total number of points in the data set. Note that when two alignments with the same search sequence are such that 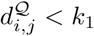, we retain only the alignment with the lowest E-value (for each query, this happens for 1% or less of the alignments).

Next, following [33] we define *γ*_*i*_ = *δ*_*i*_*ρ*_*i*_, where 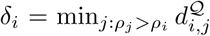, or the minimum distance of *i* to a higher density neighbour *j*. Then we sort the alignments according to decreasing values of *γ*_*i*_, Γ(*q*_0_) = {*γ* _*s*_, *γ* _*s*_ *> γ*_*s*+1_ ∀*s*}. Finally we select density peaks by identifying a *γ*_*g*_ in Γ(*q*_0_) such that 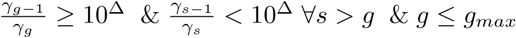. This is equivalent to looking for a gap of size Δ. between values in Γ(*q*_0_) (this was done by eyesight in [33]). We choose heuristically Δ. = 0.5 and *g*_*max*_ = 20, where *g*_*max*_ is the maximum number of peaks (primary clusters, see below) that we allow on a query sequence. As a final step, we assign to each density peak all alignments that are found at a distance smaller than *k*_1_ from the peak, and further away from any other peak: alignments mapping to a peak constitute what we call a *primary cluster*. Note that, generally, not all *B*_*i*_ alignments are assigned to a primary cluster: we discard the non-clustered alignments from the rest of the analysis.

The clusters we obtain are subsets of the previously defined 𝔅^*q*0^ ensemble, where each subset includes alignments located around the same region of the query sequence. The clustering procedure we described is schematically shown in Figure 1 (A and B), and two examples of primary clustering are shown in Figure S3.

**Figure 1:**
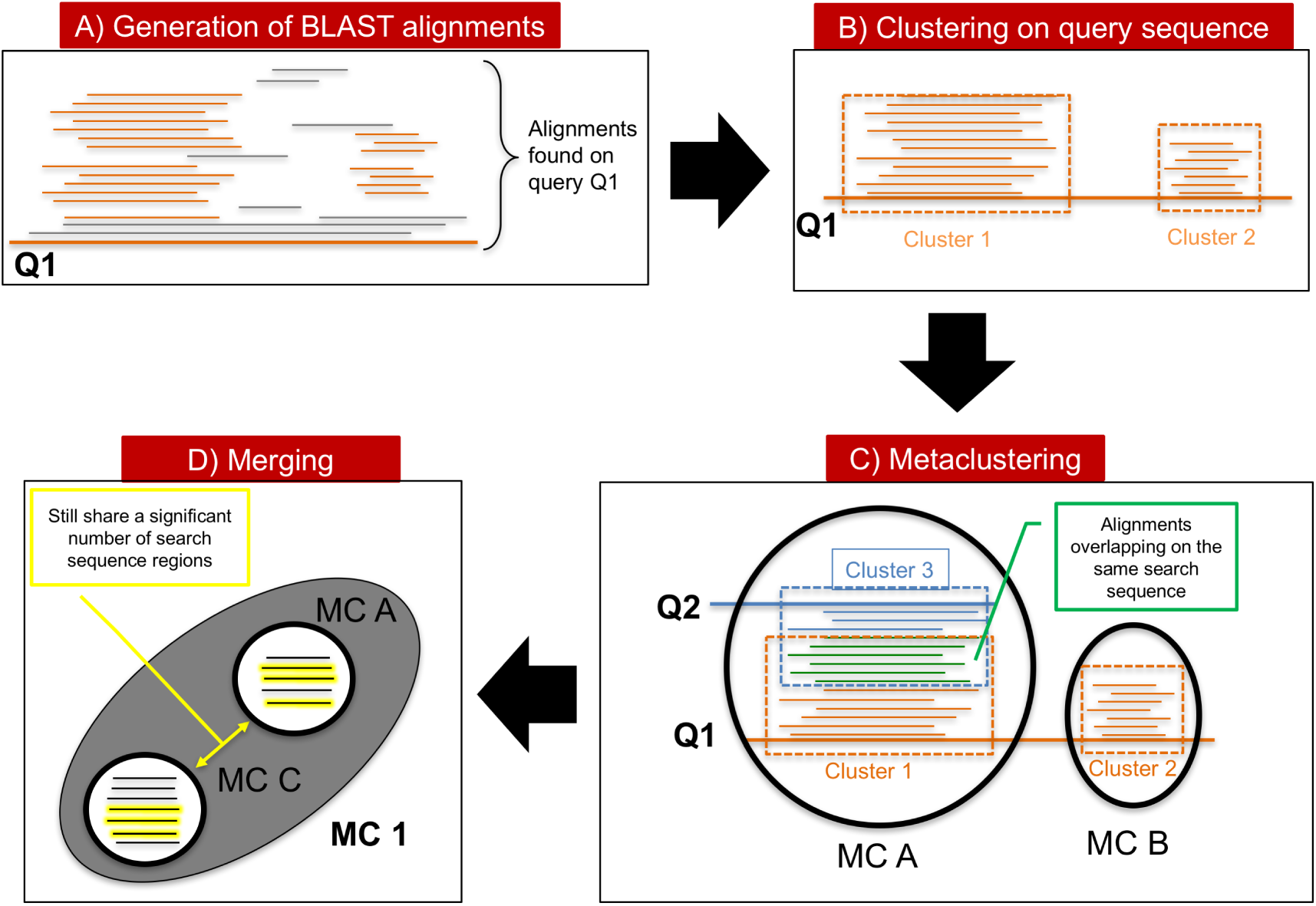
Representation of the clustering and metaclustering process. Schematically, alignments that lie on the same region of the query sequence are clustered; subsequently, different clusters are “metaclustered”, by considering close those clusters that contains a number of alignment overlapping on the same search sequence of the other cluster’s alignments; finally, metaclusters are merged by grouping those that still share a significant number of search sequence regions.

#### 2.2.2 Metaclustering

We denote the set of alignments belonging to a primary cluster *c* as 𝔅_*c*_ and we call *N*_*c*_ the number of is elements.

We define the distance between two clusters *c* and *c*_0_, associated to two queries *q* and *q*_0_ as:

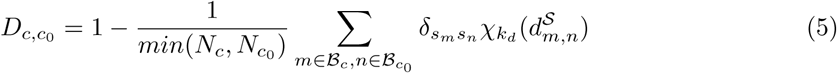

where 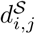 is defined as in Eq. 3 using segments *𝒮*_*i*_ and *𝒮*_*j*_ in place of *𝒬*_*i*_ and *𝒬*_*j*_, and *k*_*d*_ = 0.2 is chosen coherently with *k*_1_ in Eq.3. This distance is shorter the higher the number of alignments in the two clusters sharing the same search sequence and having a significant overlap, namely 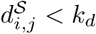.

We estimate the density *ρ*_*c*_ similarly as in Eq. 4:

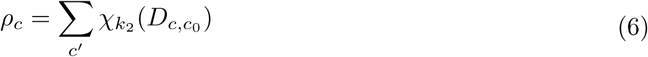

where *k*_2_ = 0.9 was also chosen following the rule of thumb in [33]. Then, similarly to what done in 2.2.1, we compute 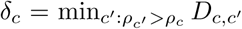. This time, however, we use a more restrictive criterion for the identification of density peaks by choosing as peaks those primary clusters for which *5*_*c*_ takes its maximum value of 1, and for which *ρ*_*c*_ *>* 1. The reason for this is that different peaks in the primary cluster space should not have significant overlaps between each other. Finally, as in 2.2.1, we assign to each density peak all primary clusters that are found at a distance smaller than *k*_2_ from the peak, and further away from any other peak; the set of primary clusters assigned to a peak constitute what we call a *metacluster* or *MC*. Primary clusters not assigned to any peak are discarded.

#### 2.2.3 Merging metaclusters

We merge similar MCs by computing the quantity 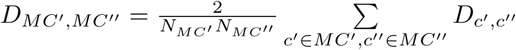 where *MC′* and *MC″* are any two metaclusters and *N*_*MC′*_ and *N*_*MC″*_ is the number of their primary clusters. *D*_*MC′, MC″*_ is the average of the distances between primary clusters contained in the two MCs. We decide to merge all MC pairs for which *D*_*MC′, MC″*_ < 0.9 based on the sorted values of all *D*_*MC′, MC″*_ obtained from clustering sequences in the PUA clan (see Support Information, Figure S2)

#### 2.2.4 Filtering metaclusters’ alignments and building profile-HMMs

A metacluster is a collection of protein regions *𝒮*_*i*_. In order to reduce the level of noise coming from outlier sequences within an MC, from the list of all regions obtained in the previous section we remove those that don’t overlap with any other sequence in the MC. More specifically, we keep region *i* if it exists another region *j* in the same MC such that 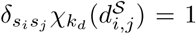. We additionally reduce redundancy at 95 percent identity using CD-HIT [22] (v4.7). MCs that at this stage contain less than 100 elements are removed from the rest of the analysis. For the sake of our comparison with Pfam annotation, only up to 5,000 members per MC are taken into consideration; if an MC has >5,000 members, we select 5,000 randomly to represent it. For the purpose of building MC-associated profile-HMMs, we further reduce MCs’ size by reducing redundancy at 60% (CD-HIT [22]) and considering maximum 1,000 members (if >1,000, we select 1,000 randomly). Next, we build an MSA using MUSCLE [9] and use the MSA to construct a profile-HMM, using HMMER (v3.1b) [26].

### 2.3 In-house Pfam annotation of the Uniref50 database and definition of Dominant Ground Truth Architectures of a metacluster

We use the Pfam classification as a gold standard to help us identifying ‘true’ evolutionary relationships between sequences in our MCs. In particular, we adopt the Pfam classification for determining both the evolutionary consistency of our MCs (homology between member regions) and the quality of their members’ boundaries. Since not all sequences in our reference Uniref50 database are annotated in Pfam, we are not able to use the Pfam database family assignments directly. Instead, we run each sequence in Uniref50 against the set of all Pfam_A.hmm models (v. 31) using the hmmscan program from the HMMER 3.1b2 suite. We assign to each protein sequence a Pfam family architecture according to the models’ manually-curated gathering thresholds. In the case of multiple significant matches overlapping along the same protein sequence, we keep only the Pfam annotation corresponding to the lowest E-value, Overlaps are calculated using start and end alignment positions; as a consequence, instances of nested domains are likely discarded (see the discussion of iron-sulfur cluster binding motifs nested within Radical_SAM domains in the Suppl. Mat.). We define the Pfam Ground Truth Architecture (GTA) *p*_*i*_ of a region *𝒮*_*i*_ as the ordered set of Pfam families that overlap with *𝒮*_*i*_, if any. The order of the families reflects their relative position along *𝒮*_*i*_. For example, suppose that we want to determine the GTA of the region of protein *s*_*i*_ =Q5BH58 spanning positions 132 to 567. Pfam annotation for Q5BH58 is as follows: PF02190 (aa 10-258), PF00004 (aa 482-625), PF05362 (aa 706-915). In this case, the GTA of *𝒮*_*i*_ is represented by *p*_*i*_ =PF02190_PF00004, comprising all Pfam families having at least a one-residue overlap with this region. We can alternatively define the GTA in terms of Pfam clans to which each Pfam family is associated (in this case *p*_*i*_(*clan*) =CL0178_CL0023); again, the GTA is an ordered string of (clan) ids. If a family is not associated to a clan in Pfam, we use the family id in the clan GTA. The boundaries of the sub-region of *𝒮*_*i*_ covered by the GTA *p*_*i*_ will be indicated as *𝒫*_*i*_. If *p*_*i*_ includes two or more families, *𝒫*_*i*_ will span also non-annotated residues between those families, if present (See Supporting Information Fig. S4). In the example above the *𝒫*_*i*_ of *𝒮*_*i*_ is the interval between residue 132 and 567. Next, we define the Pfam Dominant Ground Truth Architecture (DGTA) of a metacluster *p*^*MC*^ as the most abundant GTA (*p*_*i*_ or, alternatively, *p*^*MC*^(*clan*)) among all of its member regions. Note that the clan(s) in *p*^*MC*^(*clan*) may not necessarily correspond to the clan(s) of the families in *p*^*MC*^.

### 2.4 Comparing DPCfam metaclusters with the Pfam “ground truth”

When using Pfam annotations to analyze the evolutionary consistency of our MC classification, one should take into account the following: i) evolutionary distances between families within a Pfam clan can differ greatly; in particular, some families may be very closely related to each other. For this reason, it is often more informative to look at consistency of annotation in MCs at the clan level; ii) along with many full-length sequences, Uniref50 also contains sequence fragments. This may be relevant when comparing MC member annotations, especially for those MCs with a multi-domain DGTA. iii) Pfam classification of families and clans can be incomplete; as a consequence, regions in Uniref50 that are not currently annotated in Pfam may still belong to known Pfam families and clans.

Given an MC, we first determine its DGTAs both at the family and at the clan level and we indicate with %DAF (family) and %DAC (clan) their relative frequencies among MC members. Hereafter, we call “DGTA members” those for which, at the clan level, *p*_*i*_ = *p*^*MC*^. Next, we consider MC members that match the DGTA (again, at the clan level) only partially. While this makes sense in light of observations (ii) and (iii) above, it also allows for some variability in length among MC members. We compute the percentage of MC members with a GTA that lacks one or more of the DGTA clans but, at the same time, doesn’t feature any extra clan(s). We sum this percentage to %DAC and report it as %DACF (F=fewer); we still ask that the remaining clans are in the same order as in the DGTA. Note that MC members lacking any Pfam annotation are counted in %DACF. This is consistent with the idea that having no Pfam annotation does not imply that a region is not part of an existing Pfam clan (observation (iii) above). Finally, we compute the percentage of MC members with a GTA that features one or more Pfam clans not found in the DGTA but, at the same time, contains at least one of the original DGTA clans. We sum this to %DACF and call it %DACFA (A=additional). We will see in the Results section that the analysis of differences between these percentage scores greatly facilitates the identification of MCs that may not be evolutionarily sound, as well as, those MCs that may help improving the Pfam classification by expanding family and clan membership, by uncovering novel domains or by pointing to potential inconsistencies in the existing annotation. Comparison between the DPCfam and Pfam classifications cannot be reduced to presence or absence of families and clans on MC members. Indeed, the degree of agreement between the boundaries *𝒮*_*i*_ of the MCs’ regions and the boundaries *𝒫*_*i*_ of the Pfam annotations is also important.

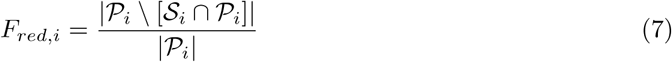

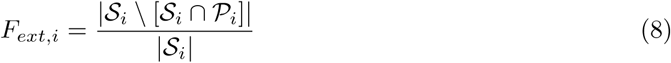

*F*_*red,i*_ represents the fraction of the DGTA *𝒫*_*i*_ that is not covered by the region *𝒮*_*i*_; vice versa, *F*_*ext,i*_ is the fraction of the region *𝒮*_*i*_ that is not covered by the DGTA. We use these two measures to characterize boundaries of entire MCs with respect to Pfam annotations by computing their average over all of the MC cluster’s DGTA members. We denote these averages as *F*_*red*_ and *F*_*ext*_.

### 2.5 Pfam clans used in this work as DPCfam query datasets

In the Pfam classification, clans group together families that are evolutionary related. As previously mentioned, families in clans may be remotely related (possibly representing domains of different function) or, sometimes, evolutionarily close (sharing a sizable number of member regions). In this work, we study two Pfam clans: PUA [15] and P53-like [4]. As of Pfam v.31, which we use throughout unless otherwise specified, the PUA clan comprised the following 11 families (25,659 sequences in total): ASCH, DUF3850, EVE, LON_substr_bdg, Methyltranf_PUA, PUA, PUA_2, TruB-C_2, TruB_C, UPF0113 and YTH. The P53-like clan, instead, comprised 9 families (8,857 sequences in total): T-Box, STAT_bind, Runt, RHD_DNA_bind, PAD_M, P53, NDT90_PhoG, LAG1-DNAbind and CEP1-DNA_bind. Both clans were selected because: 1) they are of medium size, thus rendering manual validation of results a more manageable task while still providing a rather complex set of relationships between sequences within and outside of the clan; 2) an extensive structural information is available for most of their families, which provides key insight for comparing the DPCfam and Pfam classifications. Additionally, the PUA clan is well-known to us from previous studies [5]. We note that since the DPCfam algorithm is run on full-length sequences, the MCs it generates when using PUA and P53-like annotated proteins as queries can map to any family that is found in a shared architecture with elements of those two clans.

## 3 Results

We first present in details the results obtained when running DPCfam on the PUA clan (section 3.1). Note that we used this clan to empirically adjust some aspects of our method (including parameters *k*_1_ and *k*_2_ see Eq. 4 and 6, the procedure to merge clusters and the BLAST search parameters). In the second part (3.2), we discuss clustering of the P53-like clan when using the same exact protocol used for PUA.

### 3.1 DPCfam clustering of the PUA clan

Starting from the PUA_UR50 sequence dataset (see Methods), DPCfam produces 74 MCs in total (Figure Suppl Mat. S5 for the MC size distribution). Hereafter, we focus on the subset of 32 MCs with at least 500 members (MC1_PUA to MC32_PUA, Table S1). As previously mentioned, MCs can represent single or multi-family architectures and their DGTAs can contain PUA clan families (5 in total, in bold in Table 1) or not (27 total). Also, different MCs can map to the same Pfam family or architecture.

**Table 1:**
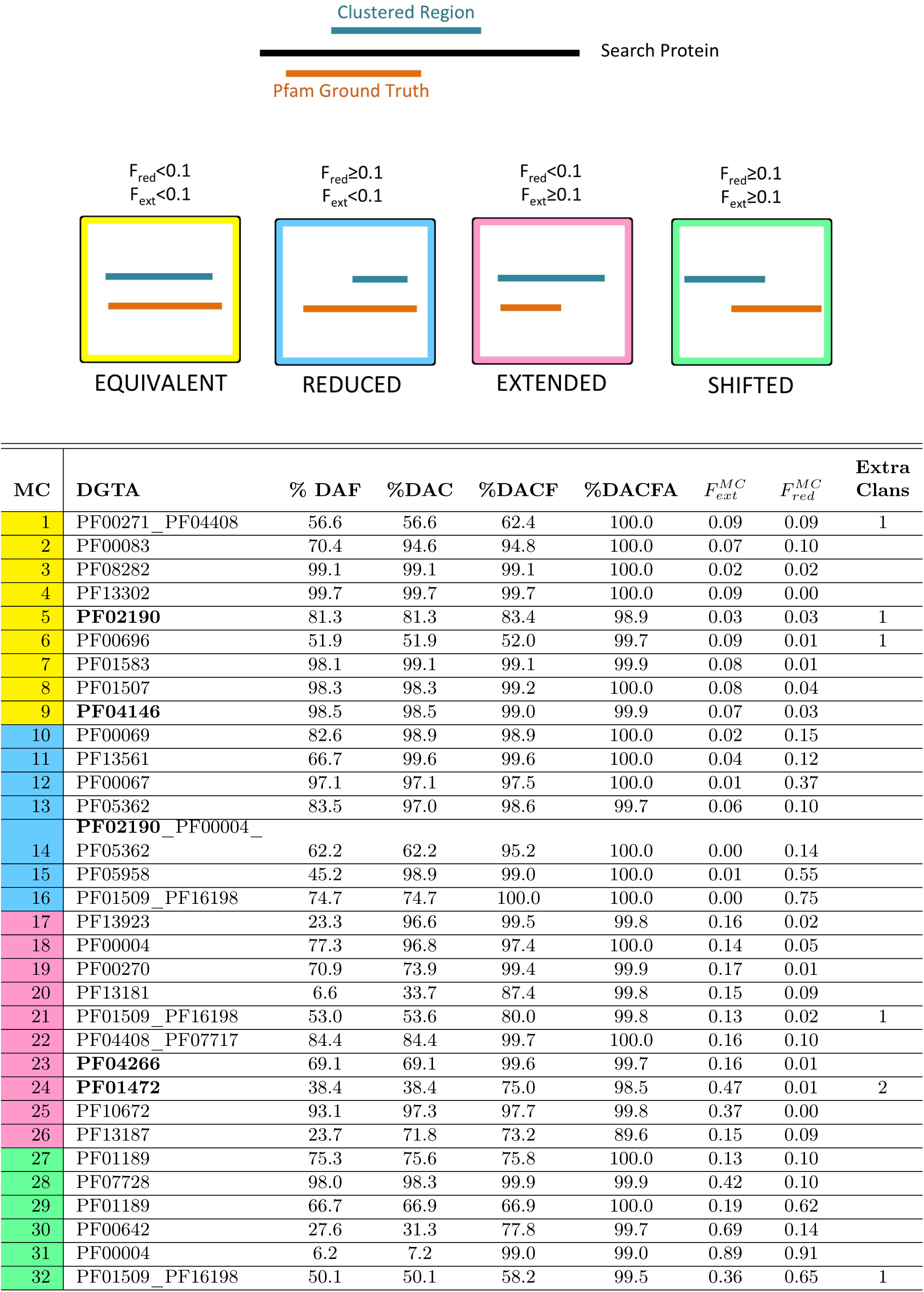
DGTA annotation of PUA_UR50 MCs,(only MCs with >500 members). Top panel: pictorial representation of how MCs are qualitatively classified based on the overlap between DGTA and DGTA members (additionally see Methods for the definition of these categories). In the Table, for each MC, we report: the family-level Pfam Dominant Ground Truth Architecture (DGTA); the percentage of members featuring a DGTA annotation either at the family (%DAF) or at the clan (%DAC) level; %DAC plus the percentage of members lacking one or more of the DGTA clans but having no additional clan’s annotation (%DACF); %DACF plus the percentage of members having clans outside of the DGTA but at least one DGTA clan (%DACFA); for DGTA members, the extent of the overlap with the DGTA, 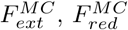; the number of extra clans that feature in %DACFA (only those present in at least 10% of clan members). MCs are colored according to a catalytic domain typical of SAM-dependent methyltransferases. MC15_PUA covers only the catalytic domain of the tRNA (Uracil-5-)-methyltransferase, albeit imperfectly (see Figure S10A). We have already discussed in previous sections several other cases with large *F*^*MC*^ values, including MC16_PUA 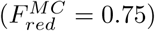 and MC24_PUA 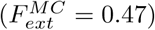, the ‘true’ DGTAs of which appear to be the single-domain architecture TruB_C_2 and the double domain-architecture pre-PUAPUA, respectively.

#### 3.1.1 Evolutionary consistency of MCs

The first question we address is whether DPCfam-generated MCs are evolutionarily consistent. In other words, we ask if MCs are formed of member sequences that share a core homologous region and can thus be used as seeds for building protein families. In Table 1, for each MC with more than 500 members, we report the percentage of member regions with a GTA that matches exactly (%DAF - family level and %DAC – clan level) or partially (%DACF and %DACFA) the DGTA of the cluster (see Methods). %DAF or %DAC close to 100% indicate that, according to Pfam, most member sequences share a homologous core region that covers all families or clans in the DGTA. For example, 99.7% of 1,795 MC4_PUA member regions are annotated in Pfam as Acetyltransf_3 (PF13302). Overall, 43.7% of MCs have %DAC>95%. Differences between %DAF and %DAC tell us to which extent member sequences are spread out across multiple families pertaining to the clan(s) represented in the DGTA. The number of Pfam families and their relative weight within an MC can be better appreciated from the graphical representation in Figure 2 (Suppl. Figure S6 for a clan level view of the same). For instance, MC17_PUA maps to several different families within the RING (CL0229) clan. This is not surprising given that the Pfam evolutionary profiles of zinc finger families within the RING clan tend to overlap (see e.g. the E-values of the families’ profile-profile alignments in the clan’s “Relationships” tab on the Pfam webserver). When we add to %DAC all those members with a GTA matching only partially the DGTA of the MC (%DACF and, finally, %DACFA) we achieve close to full coverage in most MCs. Indeed, only one MC (MC26_PUA) has %DACFA<98 (Figure 2 and, again, Table 1). Large percentage increases in the %DACF and %DACFA columns can have different meanings, but generally reflect MCs with a significant number of members sharing a core homologous region that spans less or more clans, respectively, than those found in the DGTA. Note that, in principle, it is possible for members counted as part of %DACFA to map to completely different, non-overlapping sections of the DGTA. These would be regions that are not homologous to each other. During our analysis, however, we did not come across any such outstanding example, suggesting that these are unlikely to be common occurrences. In summary, percentage values reported in Table 1 suggest that most of the MCs generated by DPCfam could be used as seeds for building homologous protein families. Another important aspect of comparing two protein classifications is to measure by how much the boundaries of the respective clusters or families differ when evaluated on the same sequences.

**Figure 2:**
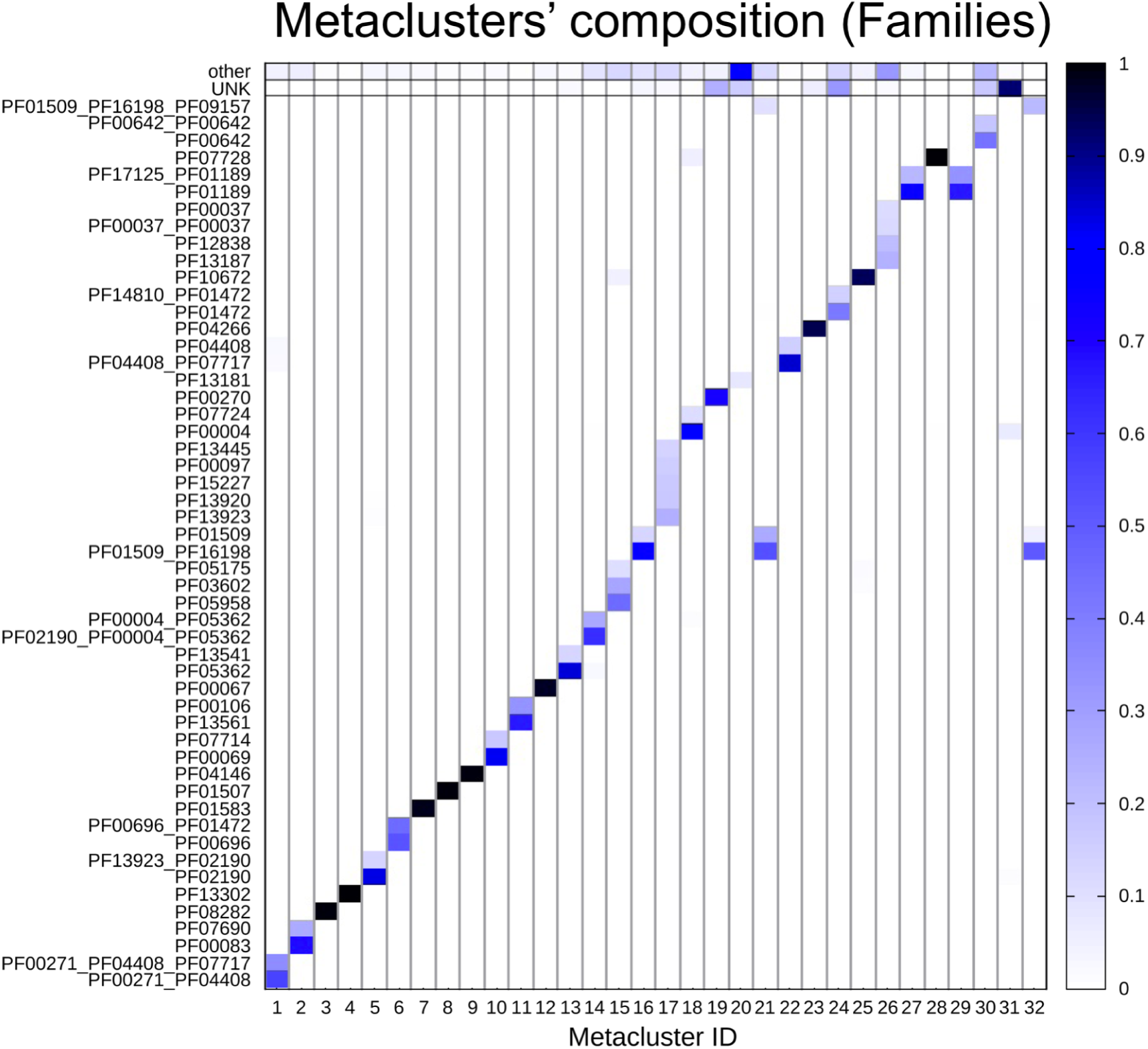
PUA_UR50 MCs vs Pfam annotation. On the x-axis, we list the MCs (sorted as in Table 1), while on the y-axis we list the Pfam GTAs (family level) represented in each MC. We report only GTAs mapping to at least 10% of MC members and we aggregate all the remaining ones under the label “other”; finally, we label “UNK” MC members with no Pfam annotation. The heatmap is colored according to GTA abundance.

The quantities 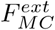 and 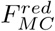 in Table 1 indicate the extent of the agreement between the boundaries of DGTA MC members and the respective Pfam annotations (these are averages over all DGTA members). To provide some qualitative insight, we classify MCs into the following four categories according to the agreement of their DGTA members with the DGTA family boundaries (see Figure in Table 1): equivalent (both 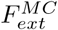 and 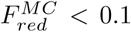, yellow; Table 1, reduced (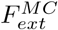 and 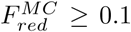, blue), extended (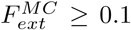 and 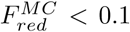, pink) and, finally, shifted (both 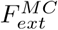 and 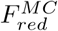 are *>* 0.1, green). We can see that the MCs in Table 1 are roughly equally split between these categories. Equivalent MCs are the closest to the DGTA architectures in terms of their boundaries; the other categories feature cases that deserve further inspection (see section 4.1.6).

#### 3.1.2 Degeneracy of MCs with respect to Pfam families

In some instances, DPCfam produces multiple clusters that map to the same Pfam family or group of families. Here it is worth pointing out that we use the DPCfam algorithm to cluster alignments rather than protein sequences. This means that alignments of the same protein region to different proteins are treated as separate entities. Our clustering protocol tries to ensure that when two regions of the same protein about the same size have a large overlap, they are classified as belonging to the same cluster. For overlaps that are small with respect to the length of the alignments being compared, the regions may end up in different MCs. One such example is represented by the trio of clusters MC16_PUA, MC21_PUA and MC32_PUA all of which feature the same DGTA, namely, TruB_N + TruB_C_2 (PF01509+PF16198). Both of these families are part of the PseudoU_synth (CL0649) Pfam clan. In Suppl. Figure S9, we show that although the 3 clusters share the same DGTA, the part of the DGTA they cover is quite different. In fact, the three MCs belongs to three different boundary categories (see Table 1): reduced (MC16_PUA), extended (MC21_PUA) and shifted (MC32_PUA). Although MC21_PUA is listed as extended, 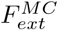 is still rather small (0.13) and the MC represents a good match to the full DGTA. MC16_PUA and MC32_PUA, instead, cover mainly TruB_C_2 and TruB_C_2 + TruB-C_2 (i.e., PUA clan family PF09157), respectively. The complex boundary relationships between these three MCs may at least in part reflect the rather complicated evolutionary make-up of the PseudoU_synth clan (see Supplementary Materials for an in depth discussion).

#### 3.1.3 MCs with potentially high levels of non-homologous member regions

Values of %DACFA significantly lower than 100% indicate the presence of a sizable subset of MC members that have a completely different clan-level Pfam annotation with respect to the DGTA. In other words, Pfam annotation for these members suggests that they may not share any region of homology with the other “DGTA-complying” MC members. From Table 1 we can see that only one metacluster, MC26_PUA, falls into this category (%DACFA=89.6). A closer inspection, however, reveals that this low %DACFA value is an artefact of our Pfam annotation protocol, which currently does not allow for domain nesting (see Suppl. Mat. for a detailed discussion of this MC, including our suggestions for improving annotation of the 4Fe-4S - CL0344 clan in Pfam). Another category of metaclusters to consider carefully are MCs that present large increases in %DACFA attributable to multiple (>1) Pfam clans not in the DGTA (last column in Table 1). This is the case for MC24_PUA. This MC has 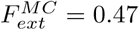, indicating that most DGTA members extend well beyond the Pfam DGTA. Additionally, a considerable fraction of members lack Pfam any annotation (37%). Most of the remaining ones feature either a TGT_C2(PF14810) + PUA architecture (14%) or a DUF1947 (PF09183) + PUA architecture (8%). Worryingly, the DUF1947 family is part of the pre-PUA clan, while TGT_C2 is currently not classified as part of a clan (that is, Pfam offers no evidence that the two are evolutionarily related). However, TGT_C2 regions are almost always found N-terminal to PUA domains and, more importantly, TGT_C2 regions display significant structural similarity to pre-PUA clan domains (see Suppl Figure S12 (Suppl Mat. MC24_PUA)). This evidence suggests that TGT_C2 is indeed likely to be a novel pre-PUA domain, thus apparently resolving the potential inconsistency we observe in the Pfam annotation of some MC24_PUA members. Interestingly, even a very sensitive profile-profile alignment method such as HHpred [38] does not appear to recognize a relationship between TGT_C2 and pre-PUA. In particular, we ran MC24_PUA MSA against Pfam v.32 and the first match to a pre-PUA clan family is represented, in 11th position, by the namesake “pre-PUA” with HHpred probability of only 27.31. This may be due to the fact that the TGT_C2 family appears to be quite divergent in sequence; for example, only a small fraction of the lone TGT_C2 region for which we have a structure (protein O58843, e.g. PDBis: 1iq8A:438-506) aligns to MC24’s profile-HMM.

#### 3.1.4 MCs with potential to extend annotation of existing Pfam families and clans

MCs with the potential to increase coverage of existing Pfam families or clans can be typically be found among those displaying large percentage increases in the %DACF or %DACFA columns. For example, metaclusters MC19_PUA and MC23_PUA feature rather large increases in %DACF (25.7% and 30.5%, respectively). Given that the DGTA of these MCs is single-domain, the increases correspond to the percentage of member regions lacking any annotation in Pfam. MC19_PUA DGTA is composed of the single helicase family DEAD (PF00270). Unannotated MC19_PUA member regions are almost always found at the N-terminus of proteins with one or more families in the Helicase_C + HA2 + OB_NTP_bind architecture. Since this is a common Pfam architecture for the DEAD domain, unannotated regions in MC19_PUA are likely to represent yet unrecognized members of the DEAD family. The DGTA of MC23_PUA, instead, corresponds to the ASCH domain (PUA clan). A lookup of MC23_PUA unannotated members in InterPro shows that many of them carry an ASCH annotation in that database, suggesting that MC23_PUA may be used to enlarge the Pfam ASCH family. In fact, MC23_PUA is particularly interesting because it appears to represent the “ASC-1 proper family” [16] potentially worth building as a separate entity in Pfam (see Suppl Mat. for more details). Similar examples are further discussed in the Suppl. Mat.

Clusters such as MC1_PUA, MC6_PUA, MC29_PUA and MC32_PUA feature sizable increases in the %DACFA column. Similar to what we saw for %DACF, a large increase in %DACFA can be ascribed to a number of reasons. Incomplete Pfam annotation of member sequences is one of them (e.g., MC1_PUA, in which several member regions are likely to lack annotation for the C-terminal domain OB_NTP_bind - PF07717). Variation in length among member regions can have a similar effect (MC6_PUA, Suppl. Mat. Figure S17). Finally, in some instances, increase in %DACFA can be due to the presence of a fraction of member regions having minimal overlap to a family outside of the DGTA. This seems to be the case, for example, of MC29_PUA with about 33% of members overlapping with a small portion of domain PF17125, located N-terminal to the DGTA.

#### 3.1.5 MCs with boundaries that differ significantly from those of the DGTA families

MCs with large values of 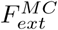 and/or 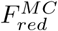, the range of which is between 0 and 1, grant further inspection. In the case of MC15_PUA, member regions typically cover only about half of the DGTA annotation on the respective full-length proteins. In particular, the DGTA is represented by the tRNA (Uracil-5-)-methyltransferase - PF05958 family. Structural data indicate that PF05958 covers two structural domains: a so-called central domain, which hosts a [Fe4S4] cluster, and

#### 3.1.6 Coverage of the PUA clan by DPCfam MCs

So far, we have looked at how consistent the Pfam annotations are within the DPCfam-generated MCs (accuracy). Clearly, it is also important to know to what extent the automatically-generated DPCfam classification recapitulates the Pfam’s one in terms of coverage of all such regions annotated by Pfam. In this section, we use all MCs (size >=100 members) created by DPCfam when run on the PUA clan and consider different values of percentage overlap between regions for calculating the MCs’ coverage of the Pfam clan. In particular, we ask how many Pfam PUA regions are covered >=25, 50, 75% or =100% by at least one MC member. In Figure S18 A, we plot the cumulative coverage of the Pfam PUA clan when ranking MCs from the one that contributes the highest coverage to the one that contributes the 15th highest coverage. We can see, for example, that >65% of PUA regions are covered for at least 75% of their length. We can ask the same question when using, instead of the MC members, the alignments generated by the profile-HMMs built from the MCs MSAs (hmmscan, E-value<10.0). This is akin to using the MCs as a source for a Pfam-style family ‘seed’ alignment. In this case, >85% of the PUA clan regions are covered for at least 75% of their length (Figure S18 B). Finally, we need to remember that most MCs cover at least some additional regions not currently annotated in Pfam that may represent new clan members (see in particular paragraph 3.1.3 above).

### 3.2 DPCfam clustering of the P53-like clan

Next, we ran DPCfam according to the same specifications described before on sequences of the P53-like (CL0073) Pfam clan and performed a similar analysis to the one done on the PUA clan. Results can be seen in Tables S2 and 2. In this case, given the smaller size of P53-like with respect to PUA, we report on all DPCfam-generated metaclusters (>100 members). Overall, the results appear to be in line with the ones obtained for PUA but for a few, interesting differences. DPCfam generates 28 MCs of size >100, of which 53.6% have %DAC>95%. Only two MCs have %DACFA<98%: MC25_P53 and MC28_P53. MC25_P53 is peculiar in that the vast majority of its members lack Pfam annotation (+95.4% in the %DACF column with respect to the single-domain DGTA). This may be explained by the high value of the low-complexity residue fraction in this MC (*LC* = 0.57, Table S2), suggesting that its member regions are unlikely to represent a structural domain. Additionally, low-complexity regions are more likely to align to non-homologous sequences (thus %DACFA<95.9%). Also in the case of MC28_P53, a large fraction of members have no Pfam annotation (54.5%); further, 33.3% are annotated as BTD (PF09270, the DGTA), 6.8% as TIG (PF01833, in a different clan with respect to BTD) and 5.3% as BTD + TIG. Large values of both 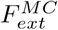 and 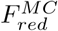 indicate that DGTA members have very little overlap with BTD and extend well beyond it. Indeed, most members cover a region C-terminal to BTD, which based on the few existing Pfam annotations and on structural considerations is likely to be a yet uncovered part of the Pfam TIG family (see Suppl. Mat. and Figure S19). Coverage of Pfam P53-like clan’s regions by MC_P53 MCs is comparable to the one observed for the PUA clan (see Figure S18). In general, in the case of P53-like, we notice two main differences with respect to the clustering of the PUA clan. First, we see a much higher degree of MC redundancy with respect to the Pfam classification. For example, 6 MCs have PF00907 as their DGTA and 4 MCs feature PF05224 in theirs. It should be noted, however, that in the case of PF00907 only two MCs have an average length of more than 50aa, which is much shorter than the length of the average protein domain [17]. In fact, MC22_P53 and MC27_P53 have length <30aa. In Figure S16 we show a graphical view of how the different MCs map to this Pfam family. Second, with respect to the PUA clan, on average, MC boundaries appear to match less well those of the DGTA families. Indeed, in Table 2 we observe several MCs with high 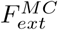 and/or 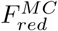.. We notice, again, that this is often the case for short MCs.

**Table 2:** DGTA annotation of P53_UR50 MCs (see Table 1 for column description.) DGTA including “(…)” represent a very long repeat, which has not been reported entirely for formatting reasons.)

| MC | DGTA | % DAF | %DAC | %DACF | %DACFA | $F_{ext}^{MC}$ | $F_{red}^{MC}$ | Extra<br>Clans |
| --- | --- | --- | --- | --- | --- | --- | --- | --- |
| 1 | PF00907 | 97.7 | 97.7 | 99.9 | 100.0 | 0.03 | 0.02 |  |
| 2 | PF00554_PF16179 | 88.8 | 88.8 | 98.1 | 100.0 | 0.03 | 0.00 |  |
| 3 | PF00400_PF00400_<br>(...)_PF00400 | 18.2 | 19.3 | 99.4 | 100.0 | 0.02 | 0.04 |  |
| 4 | PF00870 | 84.9 | 84.9 | 90.2 | 100.0 | 0.07 | 0.06 | 1 |
| 5 | PF00853 | 99.4 | 99.4 | 100.0 | 100.0 | 0.04 | 0.04 |  |
| 6 | PF00907 | 98.3 | 98.3 | 100.0 | 100.0 | 0.03 | 0.57 |  |
| 7 | PF00907 | 99.2 | 99.2 | 100.0 | 100.0 | 0.01 | 0.73 |  |
| 8 | PF00554 | 99.6 | 99.6 | 100.0 | 100.0 | 0.03 | 0.28 |  |
| 9 | PF05224 | 99.6 | 99.6 | 100.0 | 100.0 | 0.04 | 0.61 |  |
| 10 | PF02865_PF01017_<br>PF02864_PF00017 | 29.0 | 29.0 | 98.2 | 100.0 | 0.06 | 0.11 |  |
| 11 | PF00554 | 100.0 | 100.0 | 100.0 | 100.0 | 0.08 | 0.72 |  |
| 12 | PF00853 | 100.0 | 100.0 | 100.0 | 100.0 | 0.06 | 0.60 |  |
| 13 | PF09271 | 97.9 | 97.9 | 100.0 | 100.0 | 0.07 | 0.69 |  |
| 14 | PF00005 | 95.8 | 95.8 | 95.8 | 100.0 | 0.30 | 0.02 | 2 |
| 15 | PF13884 | 82.5 | 82.8 | 99.1 | 100.0 | 0.57 | 0.00 |  |
| 16 | PF05224 | 98.7 | 98.7 | 100.0 | 100.0 | 0.16 | 0.01 |  |
| 17 | PF09271_PF09270 | 86.6 | 86.6 | 97.8 | 100.0 | 0.13 | 0.01 |  |
| 18 | PF03068 | 53.0 | 53.0 | 56.6 | 100.0 | 0.12 | 0.05 | 2 |
| 19 | PF05224_PF13884_<br>PF13887 | 85.5 | 85.5 | 100.0 | 100.0 | 0.24 | 0.00 |  |
| 20 | PF09751 | 88.2 | 88.2 | 100.0 | 100.0 | 0.16 | 0.01 |  |
| 21 | PF12796_PF12796 | 26.1 | 52.2 | 68.0 | 100.0 | 0.10 | 0.11 |  |
| 22 | PF00907 | 98.1 | 98.1 | 100.0 | 100.0 | 0.13 | 0.76 |  |
| 23 | PF00907 | 99.4 | 99.4 | 100. | 100.0 | 0.14 | 0.77 |  |
| 24 | PF02864_PF00017 | 67.5 | 67.5 | 99.5 | 99.5 | 0.21 | 0.60 |  |
| 25 | PF15709 | 0.5 | 0.5 | 95.9 | 95.9 | 0.11 | 0.41 |  |
| 26 | PF05224 | 98.7 | 98.7 | 100.0 | 100.0 | 0.27 | 0.57 |  |
| 27 | PF00907 | 98.5 | 98.5 | 100.0 | 100.0 | 0.38 | 0.87 |  |
| 28 | PF09270 | 33.3 | 33.3 | 87.9 | 93.2 | 0.94 | 0.93 |  |

## 4 Discussion

Automatic classification of proteins into homologous regions or domains is a notoriously difficult problem due to the complexity of evolutionary relationships between proteins, which include but are not limited to the existence of multi-domain architectures, domain nesting and tandem repeats. Moreover, domain evolutionary divergence at the sequence level can be extremely high thus making it exceedingly difficult, if not impossible, to group into individual families all homologous regions. Finally, domain boundaries can be blurry. For these reasons, protein family databases that attempt to classify protein domains (Pfam [32], InterPro [28], CDD [24], SCOP [29], ECOD [7] to name but a few) use extensively either manual annotation or structural knowledge (often both). Nonetheless, unsupervised, automatic domain classification from sequence [14] [31] [11] is extremely relevant both to identify conserved regions that can later be manually refined and annotated to create novel families and for complementing manual classification in differential domain analysis of large datasets with a high degree of sequence novelty (such as for example sequences from environmental genomics [34] [18]. Here, we have presented a new unsupervised method based on Density Peak Clustering, which we named DPCfam, for the purpose of automatic protein domain classification. Although the clustering protocol parameters have been selected heuristically, benchmarking experiments suggest that the clustering is robust with respect to their choice and, additionally, to the size of the starting (query) dataset (Table. S3) (See Supplementary Materials). In this proof-of-principle experiment, we ran DPCfam on proteins that feature domains from two separate Pfam clans (PUA and P53-like). We showed that, in most cases, the DPCfam automatically-generated metaclusters (MCs) represent single or multi-domain architectures which, overall, display a good agreement with the Pfam annotation. In particular, we have discussed several examples of MCs mapping well to individual Pfam families. With respect to the presence of multi-domain MCs, we should emphasize that DPCfam clusters evolutionary modules (using sequence similarity) rather than directly structural domains (see definitions in [30]). Because of this, it may be difficult for our method to split into separate MCs structural domains that are only (or overwhelmingly) observed in joint architectures, unless these domains are separated by long regions of low conservation. Boundaries of MCs’ member regions show, on average, a good agreement with Pfam-defined boundaries. Further, for the two clans we have analysed, DPCfam achieves a good coverage of their member regions when considering a number of MCs that is roughly comparable to the number of families in the Pfam clans. However, we do observe especially in the analysis of the P53-like clan, a certain degree of redundancy between MCs (i.e. multiple MCs mapping to the same Pfam family). Although it is possible that this redundancy could be significanlty reduced by discarding short length MCs, especially when they are of small size, this is an area that we believe could potentially be improved upon by devising a more performant strategy for the metaclusters’ merging step.

In general, significant differences between clans exist in terms of size, evolutionary divergence, complexity of architecture and structural class of their families. Although these diversity cannot be recapitulated in full by the analysis of only two Pfam clans, it is worth pointing out that our clustering experiment did extend to numerous families outside of the PUA and P53-like clans (see both Table 1 and 2, where MCs mapping to PUA and P53-like families are highlighted in bold). This is due to the fact that DPCfam runs on full-length sequences and that about 45% and 39% of PUA and P53-like member regions, respectively, are part of multi-domain proteins.

Overall, we believe to have shown that DPCfam can support manual annotation by pointing to opportunities for expand existing families or clans and, occasionally, by identifying inconsistencies in the current classification (including incorrect domain boundaries and incomplete clan memberships). Also, we believe that DPCfam could be used effectively in combination with existing databases to expand the purpose of comparative domain analysis in datasets that include a significant fraction of unannotated sequences, such as those obtained from environmental sequencing projects. Future plans include wholesale, all vs all clustering of the Uniref50 database from which we expect to derive interesting new insights into existing family classifications and a list of potential new domains.

