## Supplementary material for "DPCfam: a new method for unsupervised protein family classification"

July 28, 2020

#### 1 Supplementary Text

##### 1.1 Additional MCs with potential to extend annotation

Three further MCs with single-family DGTA feature low %DAC and high %DACF (see Section 3.1.4 of the paper) indicating that, for the most part, they are constituted of member sequences that are devoid of any Pfam annotation; these are MC20_PUA, MC30_PUA and MC31_PUA. MC31_PUA member regions, 92% of which are unannotated, are found in ATP-dependent Lon protease proteins and typically cover a helical region located at the N- terminus of the AAA (PF00004) ATPase domain (Suppl. Fig. S8). This region could potentially be built into a short "pre-AAA" motif. MC20_PUA (54% of unannotated members) and MC30_PUA (46.5%) map, respectively, to tetratricopeptide-like repeats or TPRs (CL0020) and Cys3His zinc-binding domains (CL0537) also often found in tandem repeats. Tandem repeats such as these, which are relatively short and often feature a high degree of divergence in sequence, are notoriously difficult to classify exhaustively. It is thus not surprising that many elements of these MCs do not carry annotation in Pfam. Note that increases of %DACFA in these two MCs are mostly due to the presence of members with a higher number of repeated domains than found in the DGTA. Again, there might be scope here for using these MCs as a basis to boost coverage of the respective clans. Other families (both single- and multi-domain) display significant, if smaller, increases in %DACF, including MC14_PUA, MC16_PUA, MC21_PUA, MC22_PUA. In MC14_PUA, the main reason for this percentage increase seems related to variation in length among its member regions. In this case different Pfam annotations correspond to member regions of different length (Suppl. Mat. Fig. S17). In cases such as this, enforcement of a more uniform length composition at the MSA-building stage could be desirable.

##### 1.2 Discussion of MC26_PUA

MC26_PUA DGTA is constituted of Pfam family Fer4_9 (PF13187), which is part of the 4Fe-4S (CL0344) clan. Most families in this clan represent iron-sulfur cluster binding motifs (Fe-S BMs), often characterized by a CxxCxxCxxxC signature. PF13187 is one of the families in the clan that spans two such Fe-S BMs. Accordingly, most MC26_PUA member regions span two Fe-S BMs (alternative GTAs include, among others, the two-motif families Fer4_7 (PF12838) and Fer4_10 (PF13237), as well as, two copies of the single motif-family Fer4 (PF00037)). A fraction of members, however, are annotated as belonging to families (e.g., Radical_SAM PF04055 and DUF362 PF04015) that are found in clans other than CL0244. What is really happening in these cases, however, is that the MC members span Fe-S BM regions that are nested within these longer domains. Our Pfam annotation protocol is such that if a family matches a protein sequence starting at position A and ending at position B, the whole region between A and B is annotated as belonging to the family. In the above cases, however, the profile-HMM of the family does not match the entire region from A to B but rather has a long gap in correspondence of the Fe-S BMs (see Fig. S11 for one example). As a consequence, we can conclude that MC26_PUA member sequences consistently represent regions spanning Fe-S BMs.

#### 1.3 Suggestions on potential improvements to Pfam clan 4Fe-4S-CL0344

Analysis of the MC26_PUA metacluster lead us to look in more detail at the 4Fe-4S-CL0344 clan in Pfam; our analysis uncovered a number of potential issues concerning some of the clan's member families. In the following, we report our observations and propose changes that might help improving the clan's annotation.

The CL0344 clan groups together different types of Fe-S BMs. The clan's families are typically characterized by the presence of one or two cysteine motifs. The most common motif is constituted of four tightly clustered cysteines, that is, CxxCxxCxxxC. As mentioned a single family can cover either one (e.g. Fer4 PF00037) or two such motifs (e.g. Fer4_7 PF12838). Some outliers exhibit variations of the above cysteine signature (for example, Molybdop_Fe4S4-PF04879, FeS-PF04060 and RLI-PF04068). Some families are instead of dubious annotation. Family Fer4_24 represents a coiled-coil oligomerization domain and should hence be removed from the list of 4Fe-4S families. Similarly, family Nitr_red_alph_N-PF14710 does not cover the Fe-S BMs of the nitrate reductase proteins it is found on and should hence be removed or modified. Finally, Fer4_12 PF13353 and Fer4_14 PF13394 contain an Fe-S BM featuring 3 conserved cysteines that appears to be an integral part of the Radical_SAM-PF04055 family. In particular, PF13353 and PF13394 are not nested into Radical_SAM domain but rather overlap with the N-terminus of this family, which also covers the Fe-S BM. Additionally, and for no obvious reason, they extend well beyond the Fe-S BM, covering a sizable fraction of the Radical_SAM domain TIM barrel. In this case, if one wanted to keep these Fe-S BMs as part of the 4Fe-4S clan, PF13353 and PF13394 should be trimmed so that they include only this portion of the protein and, in order to avoid overlaps between families in different clans, the Radical_SAM family should be trimmed in the opposite direction. We note, however, that although the structural organization of the Fe-S BM of PF13353 and PF13394 is similar to the one typically observed for other motifs in the 4Fe-4S clan, profile-profile alignments (see relationships in Pfam clan) return no significant match between these two families and the other families in 4Fe-4S. This appears to suggest that if these BMs are homologous to 4Fe-4S, their homology is quite remote. As a consequence, an alternative solution could entail deleting the PF13353 and PF13394 families altogether and keep their Fe-S BMs as part of the Radical_SAM family.

#### 1.4 Discussion of MC24_PUA

MC24_PUA GDTA is represented by the PUA family (PF01472), which is the only Pfam annotation on 38.4% of MC members. The value of $F_{ext}^{MC} = 0.47$ (Table 2), however, tell us that these member regions extend well beyond the PF01472 annotation. Indeed, several members feature one additional Pfam family annotation at the N-terminus of the PUA domain: 13.6% feature a TGT_C2 (PF14810) domain, 8.4% a DUF1947 (PF09183) domain and, finally, 1% a TruB_C_2 (PF16198) domain. When present, these domains are well covered by MC24_PUA members: 97% of TGT_C2 amino acids are covered, 67% of DUF1947 and 61% of TruB_C_2. Interestingly, these 3 domains are not found in the same Pfam clan, that is, they are not recognized as homologous in the Pfam classification. DUF1947 is part of the pre-PUA (CL0668) clan that, as the name indicates, is formed of regions that are found N-terminal to the PUA domain. TruB_C_2 is part of the PseudoU_synth (CL0649) clan. TGT_C2 is not part of any Pfam clan. We first focused on the two largest sets of sequences, those containing TGT_C2 and those containing DUF1947. Structural alignment between representative structures of the two families show striking similarities (Suppl. Fig. S12) thus suggesting a common evolutionary origin notwithstanding negligible levels of sequence similarity. TGT_C2 would then be a novel pre-PUA family; this is in line with its almost exclusive association with the PUA domain in Pfam. Family TruB_C_2 is instead structurally unrelated to both DUF1947 and TGT_C2. Most of the alignments featuring TruB_C_2 have E-values of borderline significance (>0.01) supporting the notion that the inclusion of TruB_C_2 in some of the MC24_PUA members should be considered noise.

#### 1.5 Observations on the PseudoU_synth (CL0649) Pfam clan.

The PseudoU_synth (CL0649) clan has a rather complicated structure in Pfam. The domain that is covered by the clan has sometimes been split into two CL0649 families (TruB_N + TruB_C_2, PseudoU_synth_1x2, PseudoU_synth_1+DUF2344), sometimes kept as a single region (PseudoU_synth_2, TruD). The difficulty in classifying families in this clan domains comes primarily from two things: (i) the pseudouridine synthase domain appears to be formed by a tandem duplication the two moieties of which share often very little sequence similarity (and only structural similarity in terms of their general topology) and (ii) the two homologous moieties feature strand swapping and sometimes nesting of additional domains. The latter is the case for sequences in the TruD family, which in Pfam additionally covers a nested domain that should instead be built as a separate family outside of the CL0649 clan (see Fig. S13). Additionally, the boundaries of paired families such as TruB_N and TruB_C_2 don't seem to reflect the structural organization of the duplication very well (see red and blue regions in Fig. S14)). Indeed, with the current boundaries the two families represent regions of very different structure, with TruB_C_2 open and elongated structure not reminiscent of a typical structured domain. There is, for example, no pairwise structural alignment produced by DALI with default settings for the TruB_N and TruB_C_2 Pfam annotated regions of PDB structure 3u28_A. We suggest that building a family covering the entire pseudouridine synthase domain would also in this case (as in, for example, PseudoU_synth_2) be the best option. Finally, family DKCLD is not a PseudoU_synth domain and should either be included in TruB_N as a short N-terminal extension or built as a separate short motif family outside of clan CL0649 (Fig. S14)). Finally, the Pfam nomenclature of families that map to tRNA pseudouridine synthase B proteins is quite confusing. TruB_N is the N-terminal part of a PseudoU_synth domain, TruB_C is a PUA domain, TruB_C_2 is the C-terminal part of a PseudoU_synth domain and TruB-C_2 is again a PUA domain. Although we understand family names have a historical relevance, a rethinking of this particular set of names may be needed.

##### 1.6 MC23_PUA and the "ASC-1 proper family”

About 69% of MC23_PUA member regions carry an ASCH domain Pfam annotation. While the vast majority of remaining regions are not annotated in Pfam (Table 2), in InterPro many of those carry an ASCH/PUA-related annotation. Indeed, MC23_PUA is constituted of regions part of the "ASC-1 proper family”, as defined in the work by Iyer *et al.* (2005), in which ASCH domains were defined for the first time. The "ASC-1 proper family” was characterized in (Iyer *et al.* (2005)) as having a long insertion between the 3rd and 4th strand of the ASCH fold. Now that structures are available for this particular ASHC subfamily, we can additionally recognize that the domain as originally defined was cut slightly short at the C-terminus, excluding a final, extra strand (see Fig. S15). The presence of PDB structures for the C-terminal ASC-1 domain of human activating signal cointegrator 1 protein (2E5O), allowed us to build a full-length domain alignment of the family. The latter, when run against the Reference Proteomes database, appears to capture a good number of yet unannotated regions, mostly ( 40%) in Chordata.

##### 1.7 Discussion of MC28_P53

MC28_P53 contains 132 sequences, 54% of which are not annotated, 33% annotated as PF09270 (BTD), 7% annotated as PF01833 (TIG) and, finally, 5% annotated as BTD + TIG. Although BTD is not a P53-like family, it is found by the DPCfam clustering algorithm because BTD is commonly found at the C-terminus of the P53-like LAG1-DNAbind family. Although the BTD annotation is the most present in MC28_P53, the domain it represents is poorly covered by MC28_P53 member sequences. Indeed, only a few amino acids of the C-terminus of BTD as usually found in MC28_P53 members. On the contrary, when present, TIG regions are well covered. Searched with hmmsearch against the Reference Proteome dataset with MC28_P53 profile-HMM we found finding 2,083 significant hits (E-value 0.01). About half of these map to TIG domains, while the rest although often found C-terminal to a LAG1-DNAbind + BTD architecture are not annotated in Pfam. Finally, we ran MC28_P53 profile-HMM against the PDB, finding the first matches on unannotated regions of LAG1-DNAbin + BTD annotated proteins (see Figure S19 for an example). Of these, we focussed on SUH_HUMAN (Q06330), and on its structure on PDBid 3nbn_A. The region of 3nbn_A aligned to the MC28_P53's profile-HMM appears to be well-structured (in yellow in Figures S19 B and C) and is structurally similar to TIG domains (in the figure we use, in particular, PDB TIG structure 4hw6) (Figure S19 C.). In conclusion, MC28_P53 is likely to represent a TIG family, which covers a good number of TIG domains not yet annotated in Pfam.

#### 1.8 Robustness of the metaclustering procedure

We test (*a posteriori*) the robustness of the metaclustering procedure on P53_UR50 with respect to small variations (10%) of the $k_1$, $k_2$ and $\Delta$ parameters, and when reducing by half the size of the query sequence dataset. In particular, we compare the assignment of alignments to metaclusters before the filtering step (see Table S3).

In our comparison, we use: i) the number of alignments that are assigned to metaclusters; ii) the percentage of alignments metaclustered with the standard set of parameters that are still assigned to metaclusters when utilising the modified parameters; iii) the Normalized Mutual Information (NMI, see below). In general, parameters' variations do not result in significant changes in the number of alignments metaclustered as discussed below.

Variations in $k_1$ and $k_2$ imply the use of smaller or larger cutoffs in estimation of densities (see Methods 2.2.1 and 2.2.2) and a more or less restrictive criterion for assigning alignment and primary clusters, respectively, to density peaks. Not surprisingly, larger/smaller values of $k_1$ and $k_2$ translate into a higher/smaller number of metaclustered alignments (see second column of Table S3 ). This also influences the percentage of alignments in metaclusters obtained using the standard parameters that are also metaclustered with the new choices of $k_1$ and $k_2$ (see third column of Table S3). Different values of $\Delta$ lead to different numbers of density peaks (clusters) being considered and as a consequence also the number of alignments in metaclusters. Varying $\Delta$ by $\pm 10\%$ does not change significantly (by about 2%) the number of metaclustered alignments, and 98% of the alignments that were metaclustered when using the standard value of $\Delta$ are also found in the new metaclusters. When reducing the query dataset by half (we call the reduced dataset 50%_P53_UR50), we have to restrict our analysis to alignments involving the new set of queries. 644,648 of these alignments were found in metaclusters generated by the full P53_UR50 dataset, while 642,223 in metaclusters obtained from 50%_P53_UR50 and almost all of the 642,223 were in the first group (see third column of Table S3 ). Finally, we use the NMI as a quantitative measure of the consistency between metaclusters obtained with standard and alternative choices of the parameters, respectively. The NMI between two sets of clusters is defined as $\mathrm{NMI}(C_1, C_2) = \frac{2I(C_1,C_2)}{H(C_1)+H(C_2))} \in [0, 1]$, where $C_1$ and $C_2$ are labels that refer to the two alternative clustering procedures, $I$ is the Mutual Information between the two classifications and $H(C)$ is the entropy of a single classification. Two identical classifications give NMI=1. In order to compute NMI, we consider those alignments that have been metaclustered by both clustering procedures (i.e., using standard and alternative choices of parameters, respectively, third column of Table S3). NMI values are close to 1 for all comparisons (fourth column in Table S3). Taken together, these results indicate that the metaclustering procedure is robust to small variations of the $k_1$, $k_2$ and $\Delta$ parameters.

### 2 Supplementary Tables

| MC | Size | Average Length | SDL | LC fraction | MC | Size | Average Length | SDL | LC fraction |
|---|---|---|---|---|---|---|---|---|---|
| 1 | 2915 | 347.2 | *77.8 | 0.04 | 17 | 8324 | 49.1 | 5.6 | 0.00 |
| 2 | 2870 | 393.4 | 50.0 | 0.06 | 18 | 4523 | 158.3 | 36.2 | 0.02 |
| 3 | 2735 | 257.2 | 12.6 | 0.02 | 19 | 3386 | 193.2 | 17.6 | 0.05 |
| 4 | 1795 | 153.3 | 11.4 | 0.01 | 20 | 2934 | 102.9 | 12.3 | 0.02 |
| 5 | 1575 | 207.4 | 42.5 | 0.04 | 21 | 1751 | 235.9 | 25.9 | 0.02 |
| 6 | 986 | 289.8 | *57.4 | 0.05 | 22 | 700 | 259.4 | 29.3 | 0.03 |
| 7 | 851 | 164.1 | 13.9 | 0.03 | 23 | 682 | 125.1 | 16.9 | 0.01 |
| 8 | 839 | 193.3 | 26.1 | 0.01 | 24 | 675 | 148.5 | 24.0 | 0.02 |
| 9 | 791 | 152.7 | 30.5 | 0.02 | 25 | 565 | 369.6 | 35.2 | 0.04 |
| 10 | 69369 | 223.0 | 29.3 | 0.02 | 26 | 556 | 46.8 | 5.0 | 0.02 |
| 11 | 8908 | 203.2 | 19.1 | 0.05 | 27 | 3559 | 210.8 | 28.8 | 0.03 |
| 12 | 3181 | 196.9 | 21.5 | 0.02 | 28 | 1588 | 226.5 | 45.2 | 0.01 |
| 13 | 2392 | 146.9 | 40.6 | 0.03 | 29 | 1365 | 83.9 | 13.2 | 0.02 |
| 14 | 862 | 623.1 | *102.9 | 0.04 | 30 | 691 | 86.7 | 19.7 | 0.00 |
| 15 | 615 | 84.5 | 8.4 | 0.04 | 31 | 677 | 87.5 | 15.4 | 0.02 |
| 16 | 506 | 47.7 | 4.8 | 0.01 | 32 | 625 | 119.5 | 19.9 | 0.03 |

Table S1: PUA_UR50 MCs. For each MC, we report size (i.e., number of sequence members), average and standard deviation of members' lengths and, finally, the fraction of residues (of all members) that are found in low-complexity regions (LC fraction, using the segmask software included in the ncbi-blast-2.2.30+ suite (Wootton and Federhen, 1993)). We flag MCs (*) for which the SDL is larger than 50 amino acids, or about the size of a small domain.

| MC | Size | Average Length | SDL | LC fraction | MC | Size | Average Length | SDL | LC fraction |
|---|---|---|---|---|---|---|---|---|---|
| 1 (1) | 941 | 171.9 | 26.9 | 0.01 | 15 (17) | 699 | 135.1 | 21.9 | 0.02 |
| 2 (2) | 481 | 279.4 | 30.7 | 0.01 | 16 (18) | 462 | 204.5 | 31.0 | 0.02 |
| 3 (3) | 467 | 465.9 | *92.3 | 0.01 | 17 (19) | 231 | 340.8 | 52.0 | 0.05 |
| 4 (4) | 225 | 191.1 | 29.7 | 0.01 | 18 (20) | 166 | 475.7 | *107.6 | 0.02 |
| 5 (5) | 163 | 126.0 | 10.4 | 0.01 | 19 (21) | 124 | 399.5 | *58.8 | 0.03 |
| 6 (7) | 761 | 67.8 | 8.7 | 0.01 | 20 (22) | 100 | 421.4 | *53.4 | 0.05 |
| 7 (8) | 531 | 43.6 | 4.4 | 0.00 | 21 (23) | 25859 | 186.8 | 30.8 | 0.02 |
| 8 (9) | 281 | 126.1 | 13.5 | 0.01 | 22 (24) | 525 | 29.6 | 2.7 | 0.00 |
| 9 (10) | 254 | 68.2 | 7.4 | 0.01 | 23 (25) | 363 | 38.2 | 5.0 | 0.00 |
| 10 (11) | 231 | 494.3 | *106.3 | 0.02 | 24 (26) | 203 | 136.3 | 15.9 | 0.01 |
| 11 (12) | 213 | 45.8 | 2.9 | 0.00 | 25 (27) | 194 | 111.6 | 15.0 | *0.57 |
| 12 (14) | 154 | 39.0 | 3.1 | 0.02 | 26 (28) | 158 | 111.1 | 14.2 | 0.01 |
| 13 (15) | 145 | 34.0 | 3.5 | 0.01 | 27 (29) | 137 | 28.2 | 2.6 | 0.00 |
| 14 (16) | 9428 | 208.4 | 17.8 | 0.02 | 28 (30) | 132 | 137.7 | 21.0 | 0.05 |

Table S2: P53_UR50 Metacluster's properties (see Table S1).

| **Parameters** | **Number of alignments metaclustered** | **Percentage of $k_1 = 0.2$, $k_2 = 0.9$, $\Delta = 0.5$ alignments' metaclustered** | **NMI over common alignments** |
|---|---|---|---|
| $k_1 = 0.2$, $k_2 = 0.9$, $\Delta = 0.5$ | 1,350,496 | - | - |
| $k_1 = 0.2 + 10\%$ | 1,484,231 | 94% | 0.96 |
| $k_1 = 0.2 - 10\%$ | 1,351,613 | 89% | 0.99 |
| $k_2 = 0.9 + 10\%$ | 1,494,990 | 95% | 0.99 |
| $k_2 = 0.9 - 10\%$ | 1,259,045 | 93% | 0.99 |
| $\Delta = 0.5 + 10\%$ | 1,326,511 | 98% | 0.99 |
| $\Delta = 0.5 - 10\%$ | 1,374,507 | 98% | 0.95 |
| **Query Dataset** | **Number of 50% P53_UR50 alignments metaclustered** | **Percentage of 644,648 alignments metaclustered** | **NMI over common alignments** |
| P53_UR50 | 644,648 | - | - |
| 50%_P53_UR50 | 642,223 | 99% | 0.99 |

Table S3: *Metaclusters' robustness upon ±10% variation of the $k_1$, $k_2$, $\Delta$ parameters and, additionally, when reducing by half the number of query sequences.* Test are performed on the P53_UR50 dataset. We consider alignments assigned to metaclusters before the filtering step (see Methods). $k_1 = 0.2$, $k_2 = 0.9$, $\Delta = 0.5$ are the parameters used throughout the manuscript. NMI stands for Normalized Mutual Information (see Results for the definition).

#### 3 Supplementary Figures

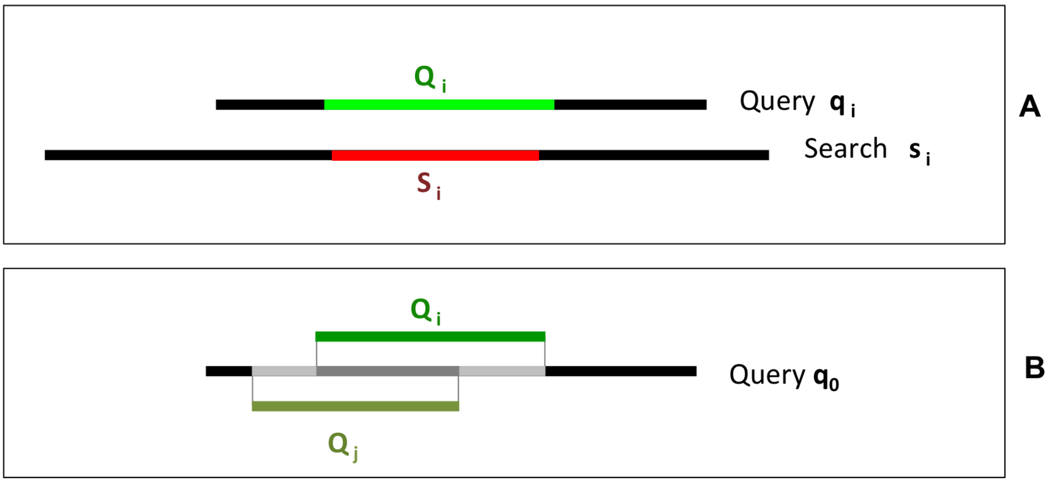


Figure S1: (A) Schematic representation of a pairwise alignment $B_i = (q_i, s_i, \mathcal{Q}_i, \mathcal{S}_i)$. The aligned regions are shown in green (query) and red (search). (B) Representation of two different alignments ($i$ and $j$) on the same query $q_0$ . The aligned regions on the query are shown in green. The dark-gray portion of the protein represents the intersection between region $\mathcal{Q}_i$ and region $\mathcal{Q}_j$ , namely $\mathcal{Q}_i \cap \mathcal{Q}_j$; ; the dark-gray+light grey region represents union of region $\mathcal{Q}_i$ and region $\mathcal{Q}_j$, namely $\mathcal{Q}_i \cup \mathcal{Q}_j$.

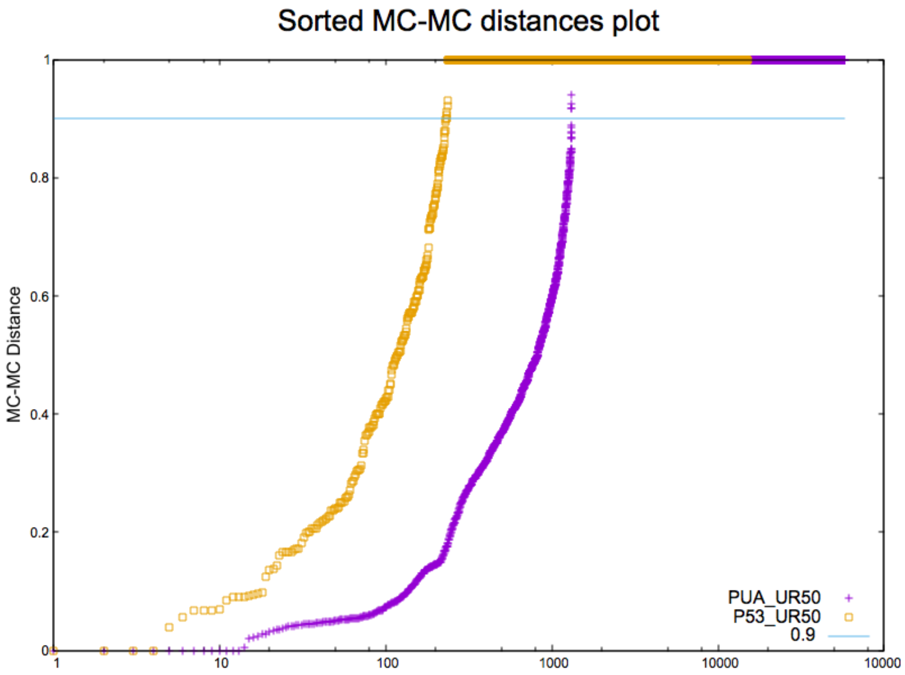


Figure S2: . Distribution of MC-MC distances for the PUA_UR50 (purple) and P53_UR50 (orange) datasets. The line corresponding to an MC-MC distance of 0.9 is our choice for the merging parameter, which has been obtained from consideration on the PUA_UR50 dataset and blindly used on the P53_UR50 dataset, with the result shown in the paper. The gap seen in the MC-MC distances of PUA_UR50 can be observed in the P53_UR50 dataset too; moreover, the curve of sorted MC-MC distances (which is strictly connected to their cumulative distribution) appears very similar. For the future work, we believe that the merging procedure should be refined, being not clear the nature of this gap and if this will be present also in other datasets.

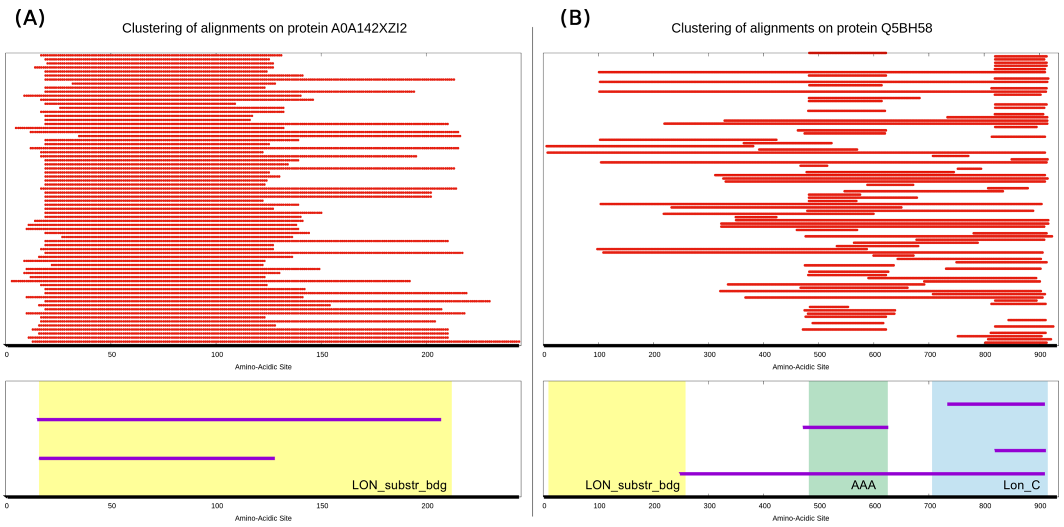


Figure S3: Examples of primary clustering results for two proteins from PUA_UR50, namely A0A142XZI2 (A) and Q5BH58 (B). Thick, black lines represent the query sequence. Red lines in the top section of each panel show a random subset of the regions of the query that have been aligned by BLAST to other sequences in the PUA_UR50 dataset. The bottom part of each panel shows instead a comparison between Pfam annotation and MC clustering of the query sequences. According to Pfam, both A0A142XZI2 and Q5BH58, contain a LON_substr_bdg domain (a member of the PUA clan), the position of which is highlighted by a yellow frame. Protein Q5BH58, in addition, contains an AAA domain and a Lon_C domain, colored green and blue, respectively. The primary clusters obtained by DPCfam using the red line alignments at the top of each panel as input are instead shown as purple lines. Primary clusters are sorted according to decreasing? value of their $\gamma$ parameter (see Methods), so that the uppers will most probably be cluster centers. We can see that some of the primary clusters overlap remarkably well with Pfam-annotated families while others either cover more than one family or overlap with only a fraction of a family. Also, note that in Q5BH58 no MC captures the LON_substr_bdg domain. In this particular case, we found that this region of Q5BH58 is a quite divergent member of the Pfam family, with both BLAST and phmmer finding less than 10 parwise alignments when using that portion of the protein as a query. Interestingly, this LON_substr_bdg region of Q5BH58 is recovered when building a profile-HMM from MC5_PUA, the DGTA of which is represented by the LON_substr_bdg domain.

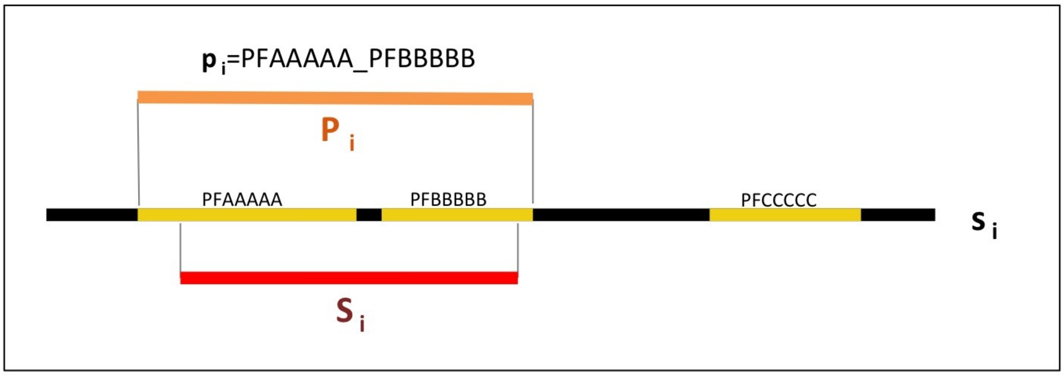


Figure S4: Schematic representation of Pfam ground truth architecture (GTA) assignment to a generic alignment $B_i(s_i, S_i)$. In this example, the full-length protein $s_i$ has the following three-family architecture: PFAAAAA + PFBBBBB + PFCCCCC; the aligned region of the search sequence, $\mathcal{S}_i$, instead covers (partially) only PFAAAAA and PFBBBBB; thus, the Pfam ground truth of $B_i$ is $p_i$=PFAAAAA_PFBBBBB (note that a 1-residue overlap of $\mathcal{S}_i$ with a family is enough for the latter to be included into the GTA); in orange we show $\mathcal{P}_i$, that is, the full region covered by the GTA families on the sequence $s_i$.

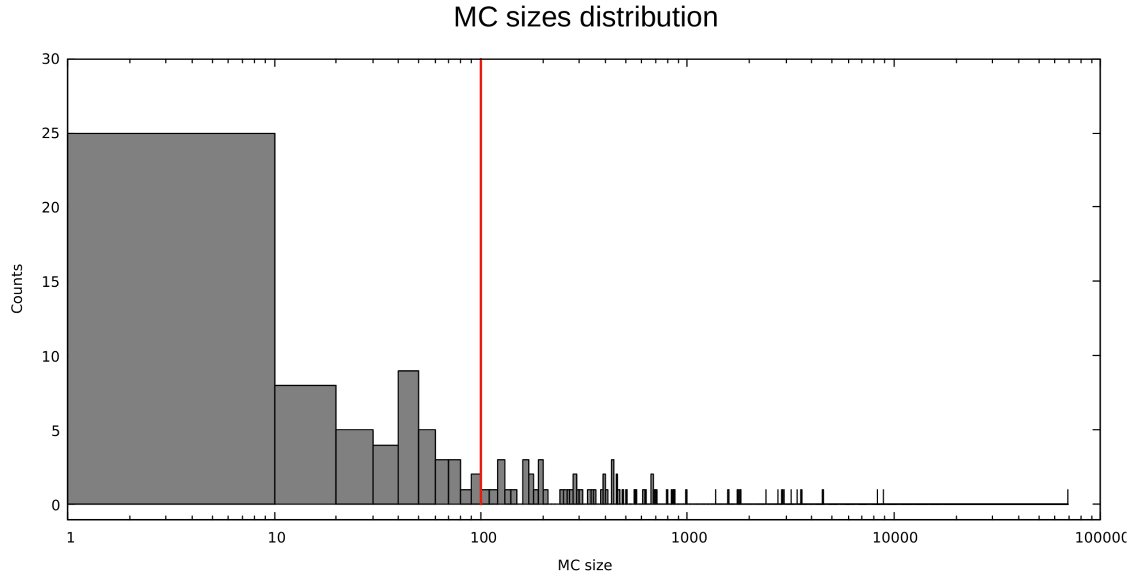


Figure S5: Size distribution of PUA_UR50 metaclusters after redundancy reduction at 95% sequence identity. Here, we include also MCs with less than 100 elements.

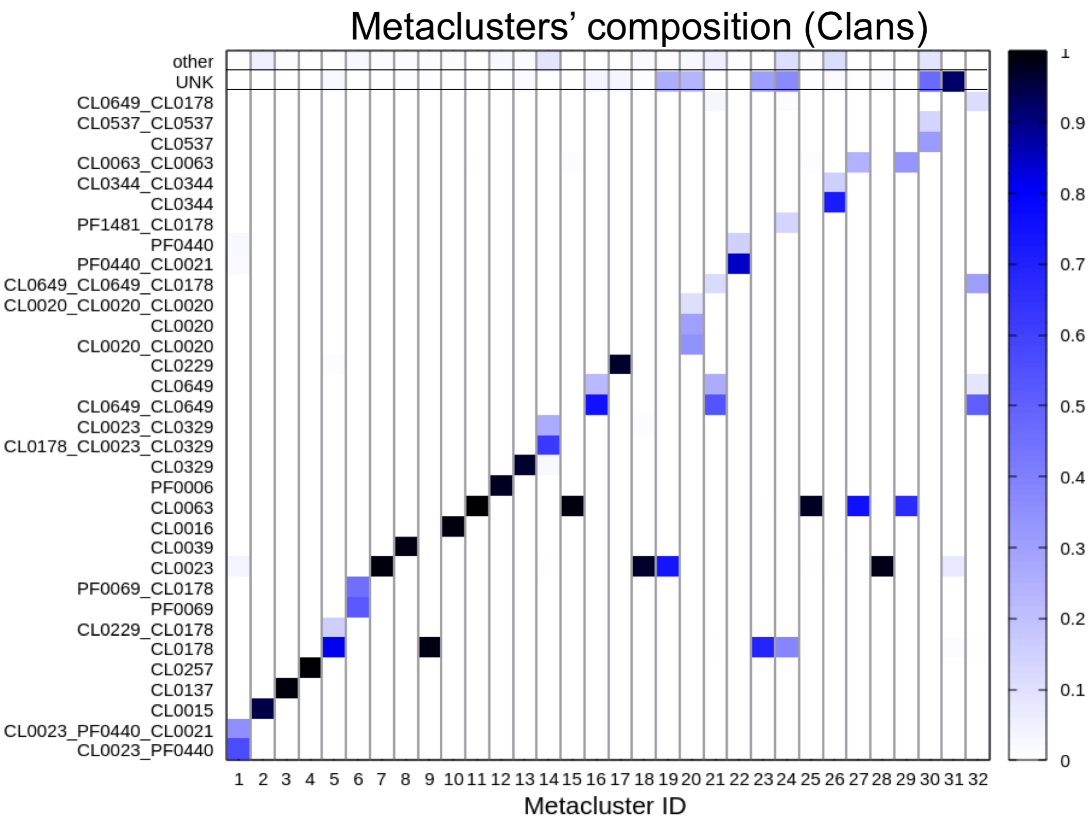


Figure S6: Density map for clan level GTAs in PUA_UR50 MCs. On the x-axis, we show the PUA MCs with more than 500 members (sorted as in Table ??), while on the y-axis we listed their GTAs. Only GTAs observed in at least 10% of members in each MC are shown, while all the remaining ones are grouped under the "other" label at the top of the map. Darker shades of blue indicate higher percentage of members with a given GTA.

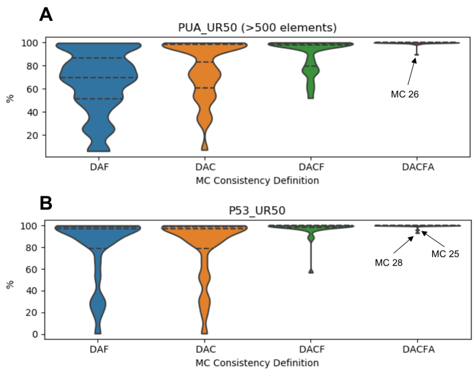


Figure S7: Violin plot of the distribution of %DAF, %DAC, %DACF and %DACFA. (A) MCs generated from the PUA_UR50 dataset (only MCs with at least 500 members) (B) MCs generated from the P53_UR50 dataset (all MCs >100 members). We label %DACFA outlier MCs, which are among the MCs discussed in the main text.

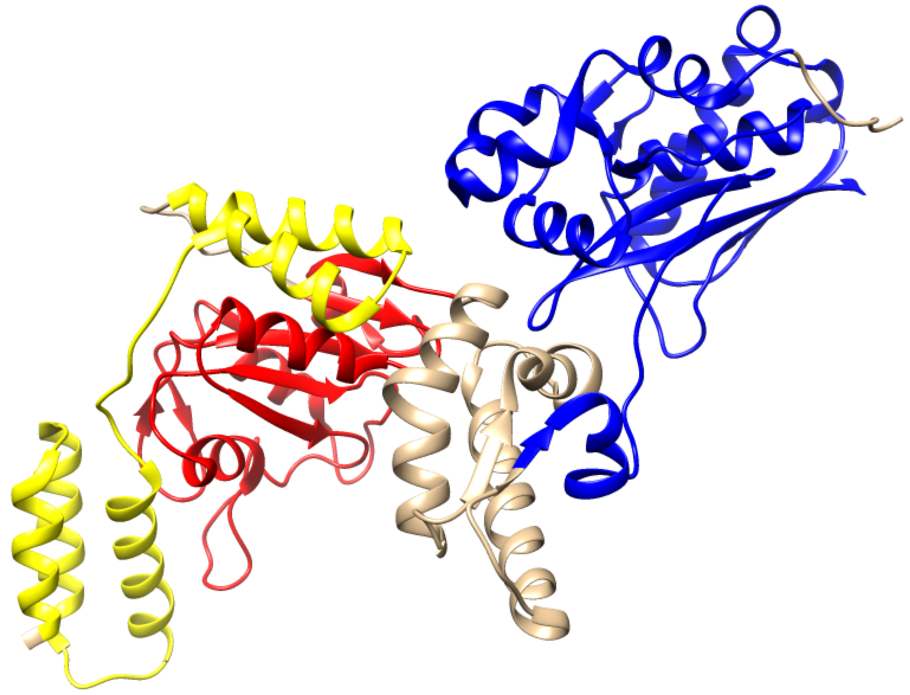

Figure S8: Structure of PDB protein chain 4ypl_A (protein A0A059VAZ3). Red and blue section show Pfam annotation: AAA region in red (aa. 351-491) and LON_C in blue (aa. 568-772). Yellow region (aa. 245-339) shows the hit of MC31_PUA profile-HMM.

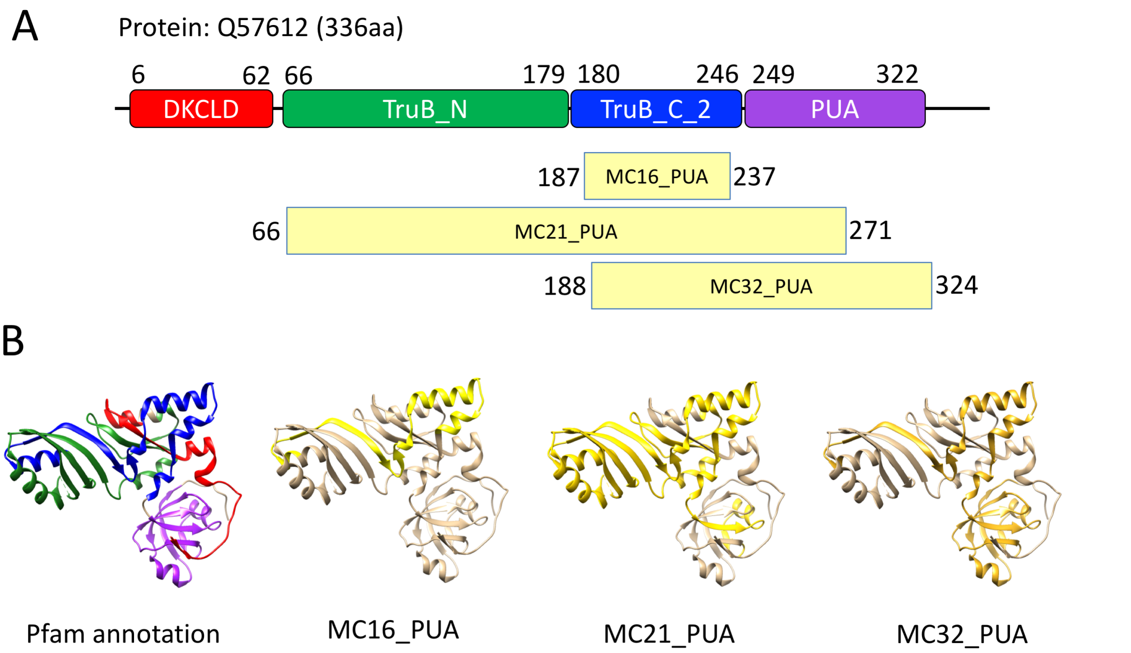


Figure S9: (A) Pfam v.32 annotation of Q57612 (protein in 2apo pdb, see panel B), containing the architecture DKCLD + TruB_N + TruB_C_2 + PUA; yellow boxes shows the regions where we find hits of profile-HMMs of MC16_PUA, MC21_PUA and MC32_PUA . (B) Structure of 2apo PDB chain A, colored following Pfam classification in panel A and according to the matches with the profile-HMMs of MC16_PUA (aa 187-237) in yellow, MC21_PUA (aa 66-271) in gold and MC32_PUA (aa 188-324) in dark gold.

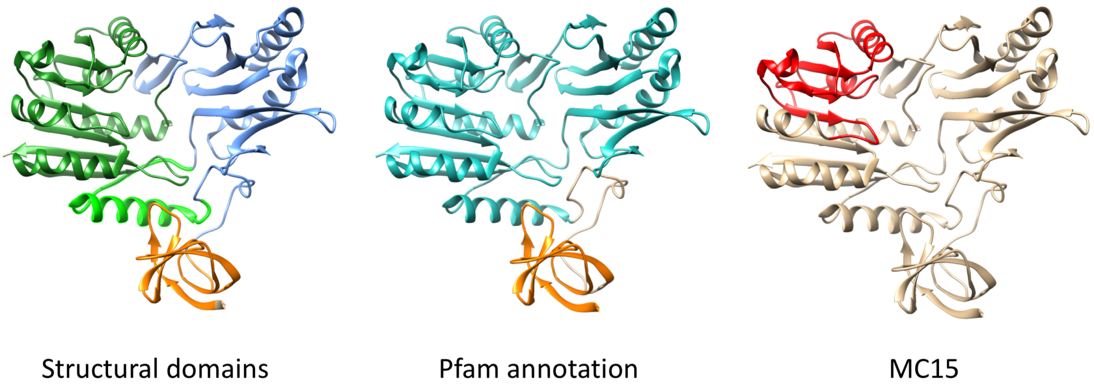


Figure S10: X-ray crystal structure of RumA, an E.coli class I SAM-dependent methyltransferase (PDB:2bh2_A). Structural domains (following (Lee et al., 2004): N-terminal domain (orange, aa15-74), Central domain (light blue, aa75–92 and 125–262) and C-terminal (catalytic) domain (light green, aa93–124 and green, aa263–431); (center) Pfam annotation: TRAM (PF01938) (cyan, aa10-67) and tRNA_U5-meth_tr (PF05958) (orange, aa95-432); (right) Region that aligns (HMMER online v3.3) to the profile-HMM built of MC15_PUA (aa285-369, red).

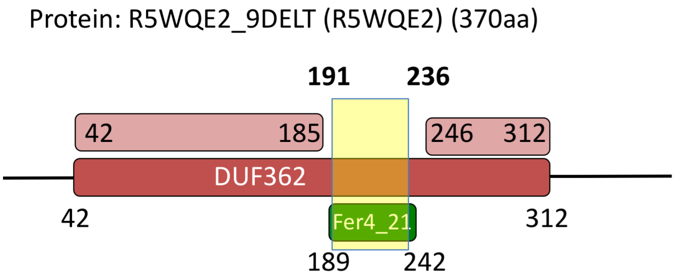


Figure S11: Example (protein R5WQE2) of nesting of an MC26_PUA region into a family of the "DUF362-like superfamily" - CL0471 clan. Solid-colored rectangles with smoothened corners mark Pfam family annotations (red and green for DUF362 and Fer4_21, respectively). The light red rectangle with smoothened corners shows the region of R5WQE2 that actually aligns to the DUF362 profile-HMM (according to hmmscan). Note that in this specific case, even in the Pfam annotation nesting of Fer4_21 into DUF362 is not accounted for. The yellow box marks the region of DPCfam MC26_PUA found on R5WQE2. Numbers represent families or region boundaries (in bold we highlight the boundaries of the MC26_PUA region). Relative lengths of boxes are approximate. Pfam annotation according to Pfam v.32.

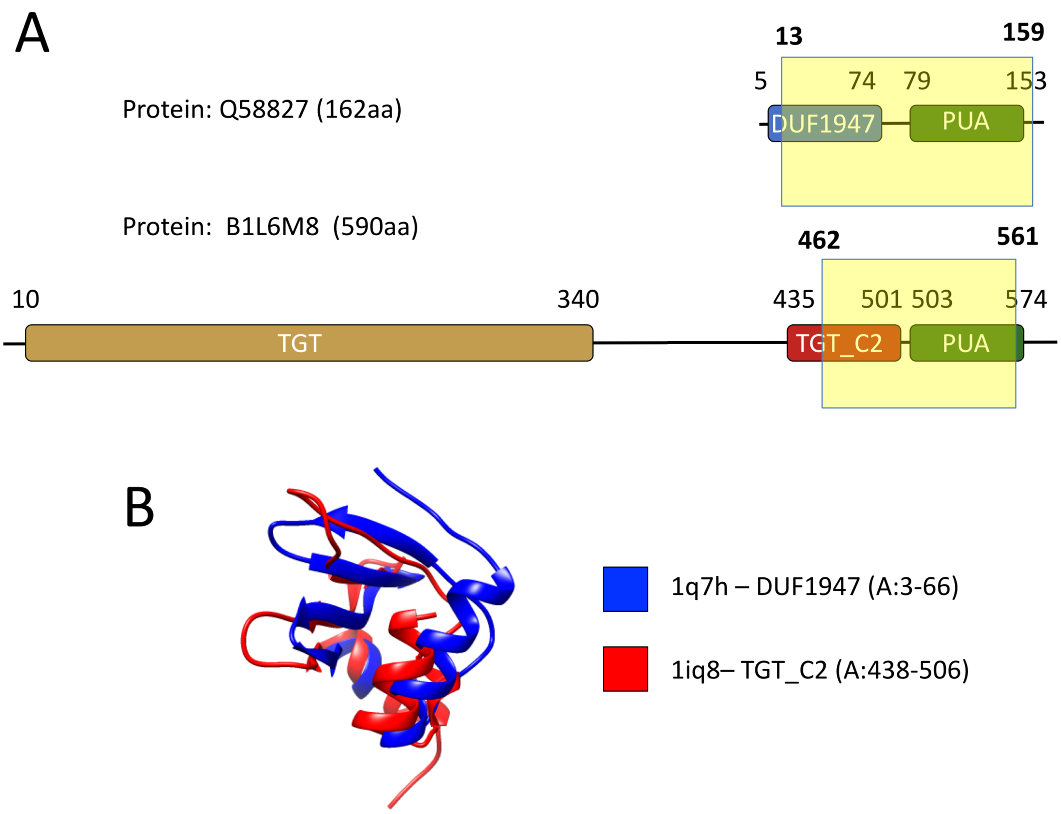


Figure S12: (A) Pfam annotation of protein Q68827, which contains pre-PUA domain DUF1947 and a PUA domain; the yellow box indicates the match MC24_PUA profile-HMM, which covers both pre-PUA and PUA domains. (B) Pfam annotation of protein B1L6M8, which contains a TGT domain, a TGT_C1, a TGT_C2 domain and a PUA domain; the yellow box indicates the match MC24_PUA profile-HMM, which covers both TGT_C2 and PUA domains. (C) Structural alignment of DUF1947 domain is pdb structure 1q7h (A:3-66) (protein Q9HIB8), with TGT_C2 domain in pdb 1iq8 (A:438-506) (protein O58843). Aligned with Dali (Holm, 2019) ; Z 4.5 , rmsd 3.0, nres 60 and %ID 12 .

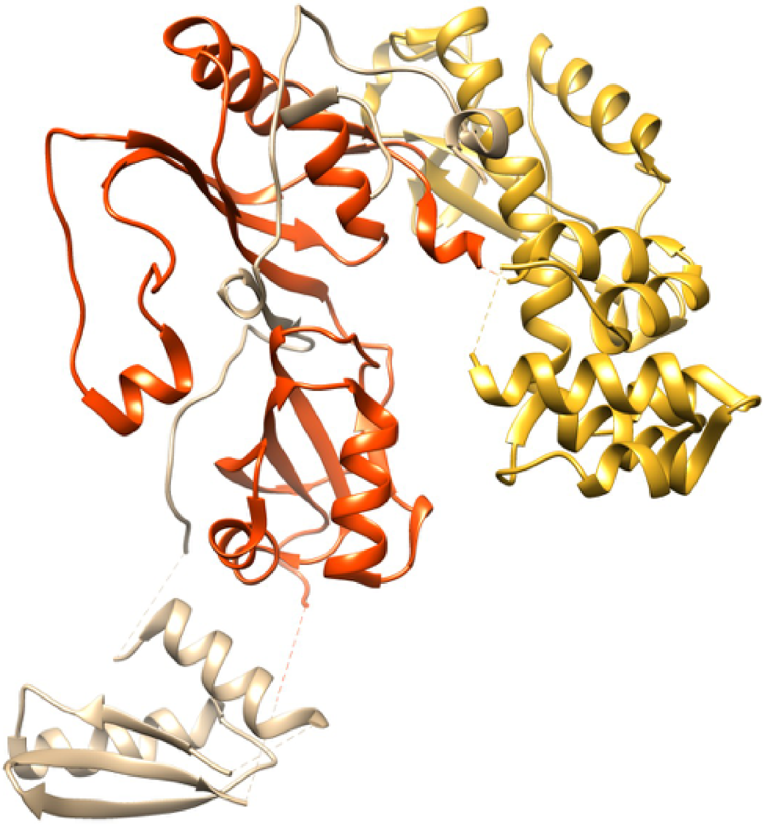

Figure S13: Structure of PDB protein chain 5kkp_A. The colored region (red+gold) covers a TruD domain as annotated in Pfam. The nested gold domain (roughly, aa384-577) is not related in structure to domains in the PseudoU_synth clan and, as such, should be built as a separate Pfam family not part of the clan. Regions not annotated in Pfam are colored tan. TruD nnotation according to Pfam 32.0.

.

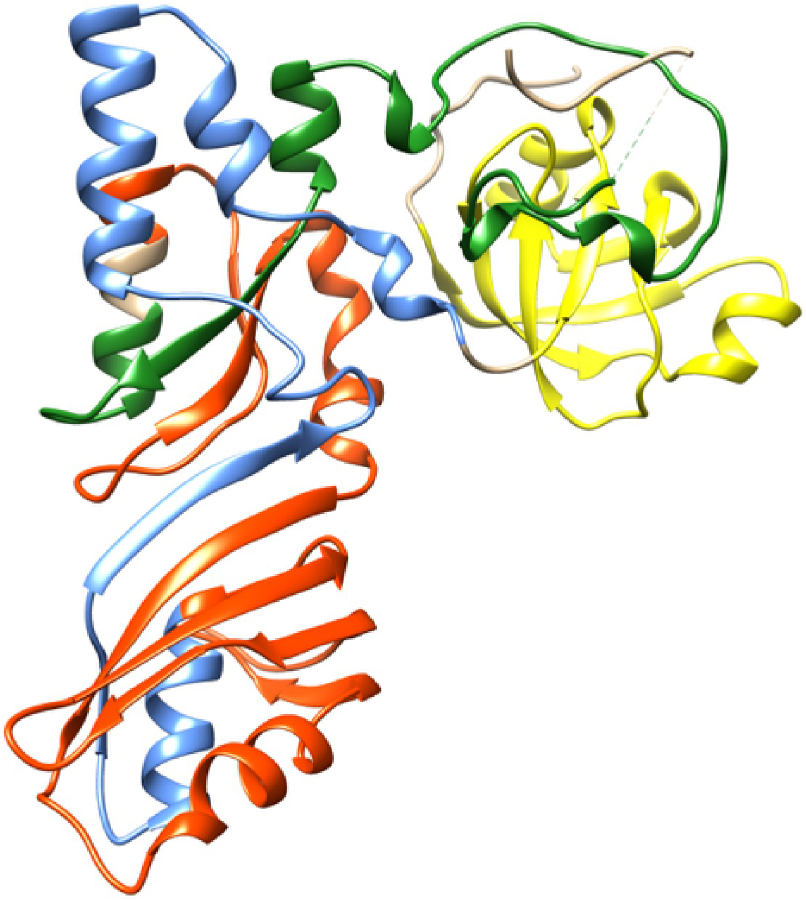

Figure S14: Structure of PDB protein chain 3u28_A. Colors identify Pfam families annotated on the structure (from N- to C-terminus): DKCLD (green), TruB_N (red), TruB_C_2 (blue) and PUA (yellow). Regions not annotated in Pfam are colored tan. Annotation according to Pfam 32.0.

.

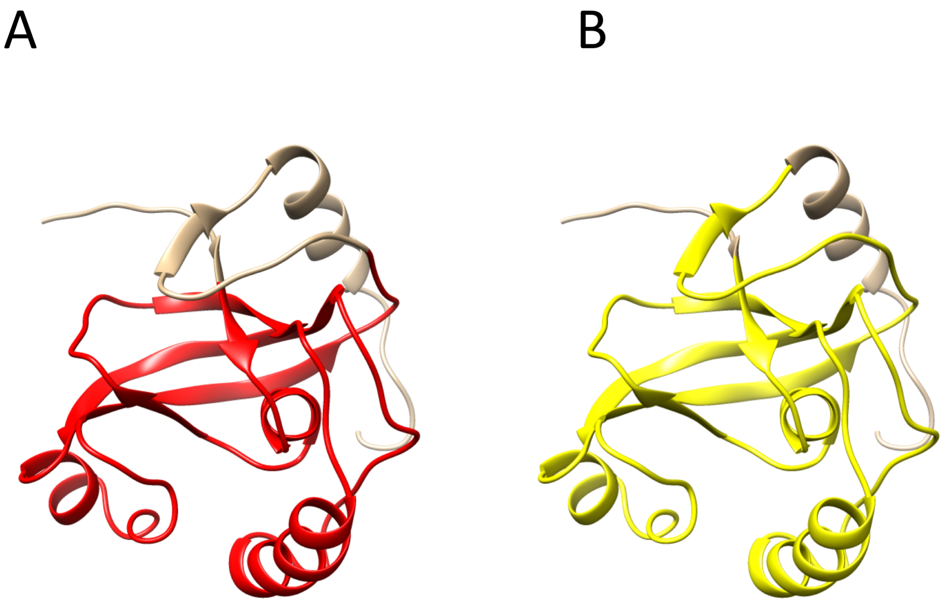


Figure S15: Structure of PDB protein chain 2e5o_A(NMR model #0.1). (A) We highlight in red the Pfam ASCH annotation, roughly corresponding to the boundaries of the "ASC-1 proper family" as found in Iyer *et al.*. It can be seen that the final strand-helix motif is missing from the annotation. (B) We highlight in yellow the region captured by MC23_PUA profile-HMM, which captures the whole ASCH region, plus the extra strand. Annotation according to Pfam v32.0.

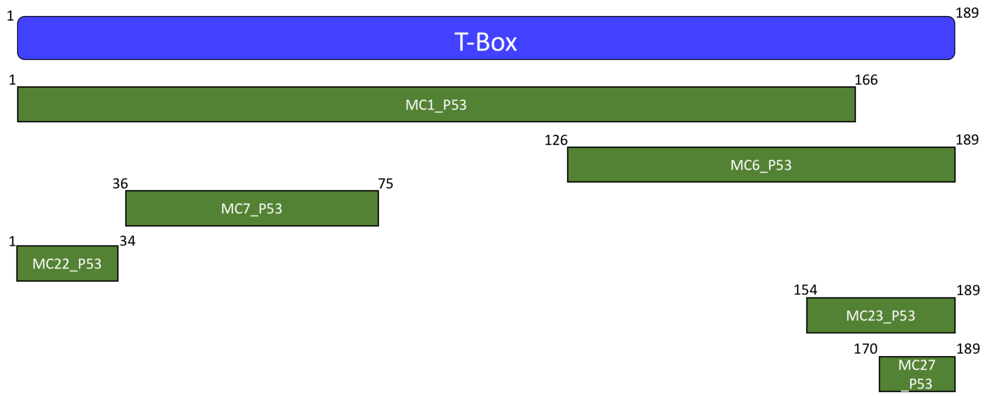


Figure S16: Coverage of P53_UR50 redundant metaclusters with respect to their common PF00907 (T-Box) DGTA. We used HHpred (Soding *et al.*, 2005)(Zimmermann *et al.*, 2018) to determine the position of each MC with respect to the T-box profile-HMM (the first match for all these MCs, with hhpred probability>98%).

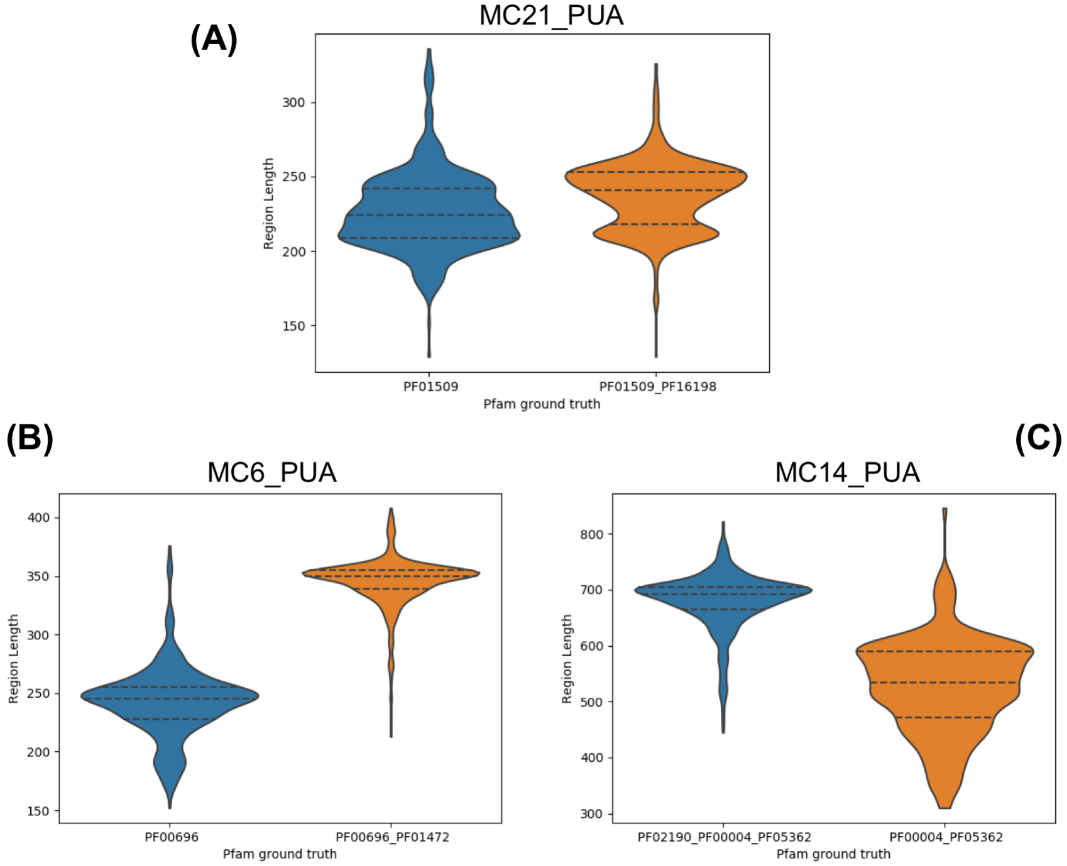


Figure S17: Distribution of member regions' length for MC21_PUA (A), MC6_PUA (B) and MC14_PUA (C). For each plot, we show the distribution of lengths of DGTA regions (i.e. matching the DGTA exactly) (blue) and, additionally, of those matching the second most abundant Pfam ground truth in the MC (orange). The different distributions observed suggest that in the case of MC21_PUA, the absence of the PF16198 family from the DGTA is likely due to a incomplete annotation of this domain in Pfam, while in the case of both MC6_PUA (B) and MC14_PUA (C), the MCs are constituted of groups of members of different length and hence as a consequence feature different, if overlapping, Pfam annotations. MC6_PUA (B) and MC14_PUA (C) length inconsistencies can be easily resolved, for example, by trimming the respective MSA alignments and removing N- and C-terminal columns with a percentage of aligned sequences $<50\%$ of the total.

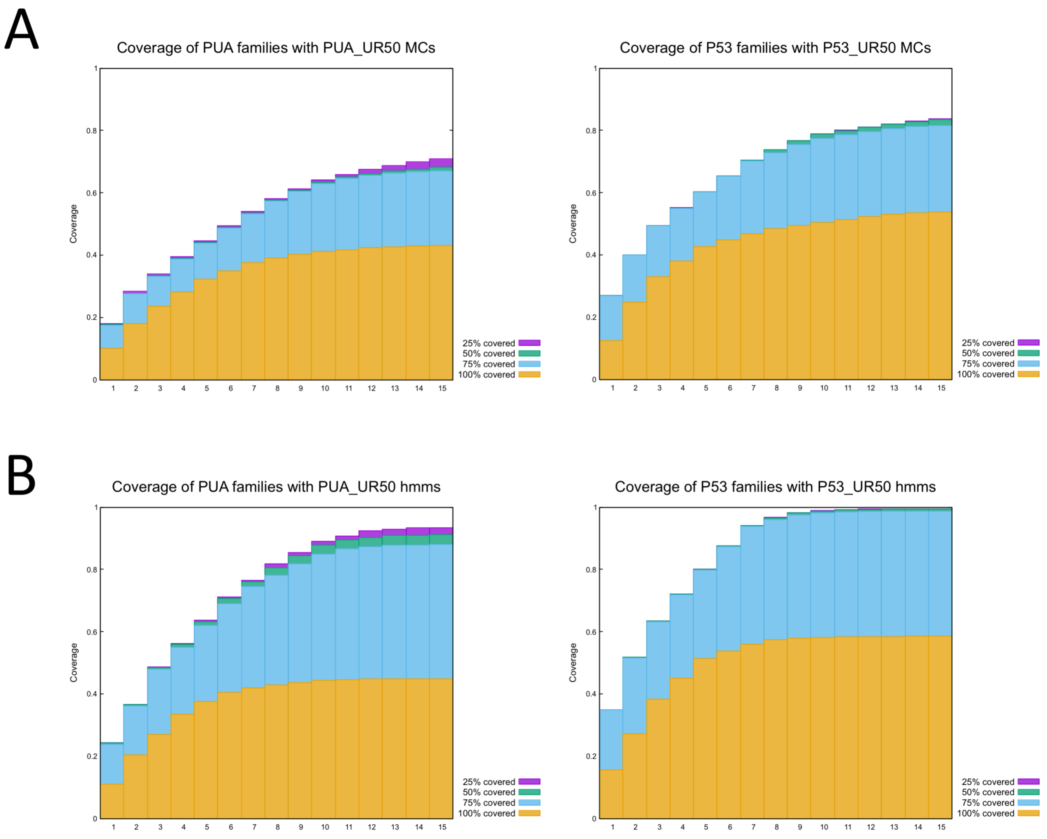


Figure S18: Coverage of PUA and P53-like clan regions. We extracted PUA and P53-like regions from Uniref50 (v.2017/7) using PfamA profile-HMMs (v.31) of the clan's families. (A) We checked if PUA (clan) regions where captured by PUA_UR50 MCs, and P53-like (clan) regions by P53_UR50 MCs, using four coverage criteria: >=25, 50, 75% or =100% of the Pfam region is covered by at least one MC member region. The two graphs gives the cumulative distribution of the respective coverage criterion for the two datasets, starting from the best-scoring MC. (B) Same as A, but with profile-HMMs of PUA_UR50 and P53_UR50 (see methods)

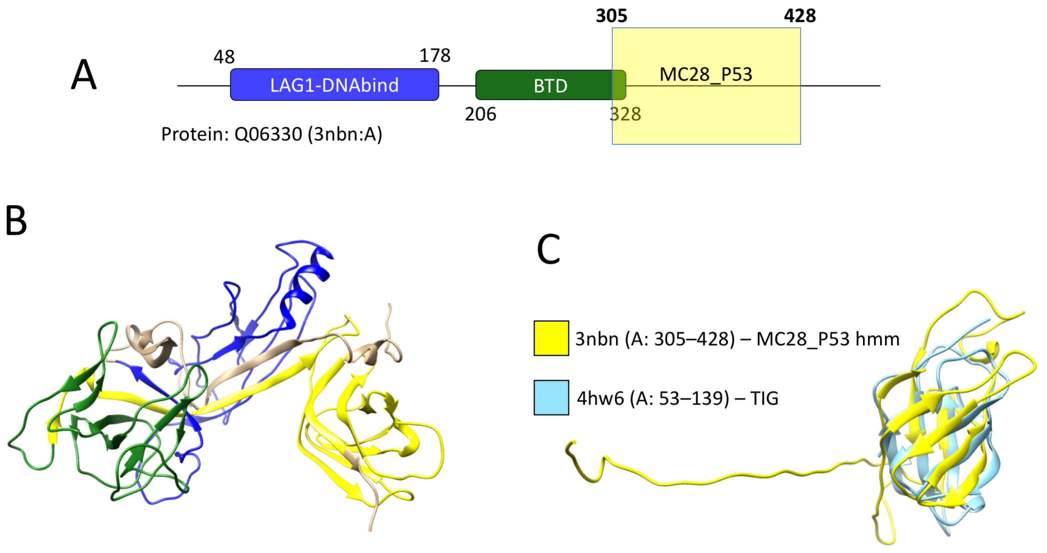


Figure S19: MC28_P53. (A) Pfam annotation for protein Q06330. Solid-colored rectangles with smoothened corners mark Pfam families (blue for LAG1-DNAbind, green for BTD). The yellow box marks the DPCfam MC28_P53 region found on the proteins running MC28_P53's profile-HMM against the PDB. Numbers represent families or region boundaries (in bold we highlight the boundaries of the MC24_PUA region). Relative lengths of boxes are approximate. Pfam annotation according to Pfam version 32.0. (B) Pfam and MC28_P53 annotations of panel (A) mapped to one of the available structures of Q06330 (PDBid 3nbn:A). Color code for fmailies and regions is the same as in (A). (C) Structural alignment between the MC28_P53's annotated region of 3nbn (yellow) and the TIG domain of PDB structure 4hw6 (light blue). Alignment obtained with DALI pairwise online tool; alignment features: Z=6.2, RMSD=2.2, percent sequence identity=25).
